# Polyglutamine-expanded TBP rewires neuronal and muscle proteomes beyond transcription in a *Drosophila* model of SCA17

**DOI:** 10.64898/2026.10.20.760399

**Authors:** Matthew V. Prifti, Kristin Richardson, Fatimah H. Hussein, Truman Kula, Anna Stephan, Yong-Dong Wang, Wei Wang, Anthony A. High, Suresh Poudel, Fabio Demontis, Sokol V. Todi

**Affiliations:** Department of Pharmacology, Wayne State University School of Medicine, Detroit, MI 48201, USA; Department of Developmental Neurobiology, St. Jude Children’s Research Hospital, Memphis, TN 38105, USA; Department of Cell and Molecular Biology, St. Jude Children’s Research Hospital, Memphis, TN 38105, USA; Center for Proteomics and Metabolomics, St. Jude Children’s Research Hospital, Memphis, TN 38105, USA; Department of Immunology, St. Jude Children’s Research Hospital, Memphis, TN 38105, USA; Department of Neurology, Wayne State University School of Medicine, Detroit MI 48201, USA

**Keywords:** Ataxia, Muscle, Neurodegeneration, Post-translational modification, Proteomics

## Abstract

Spinocerebellar ataxia type 17 (SCA17) is an age-related neurodegenerative disorder caused by polyglutamine expansion in the TATA-binding protein (TBP). TBP is a core transcription factor, but pathogenic polyglutamine expansion likely rewires cellular programs beyond transcription. Here, we utilized *Drosophila* strains that express wild-type (WT) or pathogenic (SCA17) TBP in neurons or skeletal muscle, which we analyzed vs. controls via TMT mass spectrometry, profiling of eight post-translational modifications, and RNA-seq. Shared TBP-WT and TBP-SCA17 responses predominated, impacting ribosomes and mitochondria while repressing innate immunity. Disease-specific effects were narrower and tissue-dependent: SCA17 activated xenobiotic, retinoid, and glutathione-redox pathways in the CNS, but attenuated anabolism in muscle compared to TBP-WT. Many proteomic changes occurred independently of mRNA changes, implicating post-transcriptional mechanisms. Knockdown of select SCA17-regulated proteins reduced cytotoxicity, validating omics-identified modifiers of SCA17 pathology. These findings distinguish shared TBP dosage effects from disease-linked, tissue-specific polyglutamine responses and identify evolutionarily conserved targets for SCA17.

## INTRODUCTION

Spinocerebellar ataxia type 17 (SCA17) is a neurodegenerative disorder caused by expansion of a polyglutamine (polyQ) tract within the TATA-binding protein (TBP), an essential factor for RNA polymerase II transcription initiation^1^. PolyQ disorders form a larger genetic family driven by CAG repeat expansion in coding regions, producing proteins with abnormally long polyQ tracts. The nine canonical members include Huntington’s disease (caused by polyQ expansion in HTT), dentatorubral–pallidoluysian atrophy (ATN1), spinal-bulbar muscular atrophy (AR), and several spinocerebellar ataxias (SCA1/ATXN1, SCA2/ATXN2, SCA3/ATXN3, SCA6/CACNA1A, SCA7/ATXN7, and the focus of this work, SCA17/TBP)^2,3^. Despite a shared mutational mechanism, these disorders exhibit cell-type selectivity, implying that each expanded protein interacts with specific partners and tissue-specific biological programs.

In SCA17, this selectivity is especially paradoxical because TBP is ubiquitously expressed and participates broadly in transcription initiation^1,4–6^. A central unresolved question is, therefore, how a global perturbation of TBP produces molecular responses that differ between vulnerable neural and peripheral tissues. This distinction is therapeutically important because interventions that alter TBP abundance, aggregation propensity, protein quality control, or downstream response pathways may have different effects across diverse tissues and disease stages^7–14^.

Prior SCA17 studies identified discrete mechanisms involving altered transcription-factor interactions, impaired chaperone and endoplasmic-reticulum (ER) stress responses, calpain activation, and muscle structural defects^5,6,12,15–18^. Systems-level analyses have been informative but have generally examined isolated cells, whole animals, or individual tissues and disease stages. Moreover, the design of these studies typically did not include an analysis of wild-type TBP, therefore leaving unresolved whether the observed cellular and molecular changes were indeed due to TBP polyQ expansion or, alternatively, to increased TBP expression. This distinction is critical because wild-type human TBP is itself phenotypically disruptive in the matched fly system^19^. A mutant analysis without a wild-type TBP control may therefore misclassify effects induced by TBP expression as effects caused by TBP-polyQ pathology. Finally, neural and muscle compartments have not been compared under matched genetic, environmental, temporal, and analytical conditions, and the concordance among transcript abundance, protein abundance, and post-translational modification (PTM) remodeling that occurs in the early phase of disease etiology before overt dysfunction remains unexplored. Resolving these gaps in attributions, tissue, disease-stage, and regulatory layers is necessary to distinguish between responses caused by TBP dosage vs. those that are polyQ-dependent and hence pathogenic, and the tissue-selective liabilities to TBP-polyQ that may require distinct therapeutic strategies.

*Drosophila melanogaster* offers an efficient and tractable platform for addressing these outstanding questions^20–23^. First, TBP is highly conserved between flies and humans, supporting cross-species inference of core mechanisms^15,19,24–34^. Second, its genetic tools allow precise manipulation of TBP expression in defined cell types and with high replicate numbers. Moreover, the rapid sampling of heads (enriched for neurons and glia) and thoraces (enriched for indirect and direct flight muscles and associated motor circuitry) can be used to compare TBP-mediated proteomic remodeling in neuron-rich vs. skeletal muscle-rich contexts under identical environmental and genetic conditions, thus controlling for confounders that often complicate human and mammalian studies.

In this study, we utilized a fly SCA17 toolkit based on transgenic lines that express full-length, HA-tagged human TBP with either a wild-type polyQ length (Q25; “TBP-WT”) or a patient-range expansion (Q63; “TBP-SCA17”). Both constructs were site-specifically integrated at the same genomic landing site, attP2, using the phiC31 integrase, thus minimizing position effects and ensuring comparable expression levels across these distinct transgenic lines. We previously found that ubiquitous induction of the normal or pathogenic version of human TBP had deleterious consequences in developing flies and impacted the longevity and motility of adult escapees (^19^; figure 1A). In adults, TBP-SCA17 expression produces an age-progressive accumulation of SDS-resistant species relative to TBP-WT, while both TBP-WT and TBP-SCA17 reduce lifespan and mobility in a tissue-dependent manner (^19^; figure 1A). Tissue-specific drivers further demonstrate that neuronal, glial and retina-specific expression each elicit robust phenotypes, and that polyQ-expanded TBP is generally more toxic than TBP-WT (^19^; figure 1A). Collectively, this extensive validation indicates that this model is well suited to disentangle dosage-vs.-mutation effects of TBP at the whole transcriptome and proteome scales.

**Figure 1:**
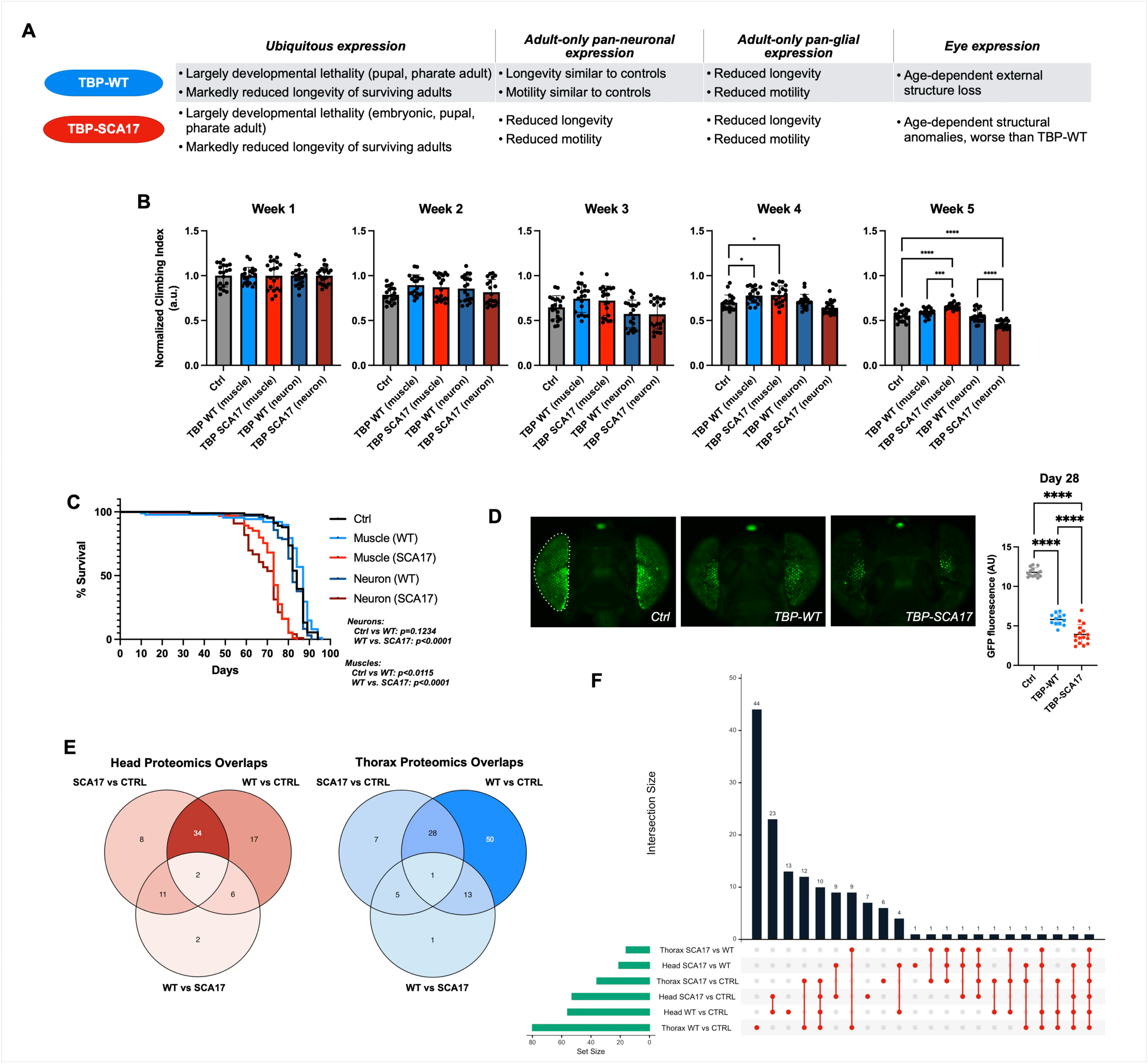
Effects of expression of normal and pathogenic TBP in *Drosophila* tissues. A) Summary of findings from prior work with the SCA17 lines ^19^. B) Effect of TBP-WT and TBP-SCA17 on fly mobility when expressed in the indicated tissues throughout development and in the adult stage. N=100 per cohort. Statistics: one-way ANOVA with Tukey’s post-hoc test. *P<0.05; **P<0.01; ****P<0.0001. C) Effect of TBP-WT and TBP-SCA17 on fly longevity when expressed in the indicated tissues throughout development and in the adult stage. N=91 for CTRL, 98 for Mef2/WT, 94 for MEf2/SCA17, 98 for elav/WT, 98 for elav/SCA17. Statistics: log-rank tests. P values are noted in the figure. D) Effect of TBP-WT and TBP-SCA17 expression in the fly retina, using CD8-GFP fluorescence as readout. At least 13 fly heads per genotype were assessed. Statistics: Brown-Forsythe and Welch’s ANOVA tests. ****P<0.0001. Left to right on the graph: N=14, N=12, N=16. E) Venn diagrams summarizing similarities and differences among genotypes and tissues. Supplemental data S1 provides a detailed list of this information. F) UpSet plot that quantifies the intersections across the six comparisons in the two tissues.

Here, we leveraged these models and used a three-genotype, tissue-resolved design to separate the broad molecular responses caused by exogenous TBP expression from those caused by polyQ expansion. Driver-only controls and matched lines expressing TBP-WT or TBP-SCA17 were analyzed in heads (enriched for central nervous system, CNS) and in thoraces (enriched for skeletal muscles). We integrated 18-plex TMT (tandem mass tag) mass spectrometry, JUMP-ptm profiling of 8 post-translational modifications (PTMs), and RNA-seq to define changes shared between normal and expanded TBP expression (i.e., TBP-WT or TBP-SCA17 vs. controls) and, conversely, to identify polyQ-specific responses (divergent from those induced by TBP WT) within each tissue. We further cross-compared these integrated omics to assess concordance across the levels of mRNAs, proteins, and associated PTMs. Through functional testing of selected candidates, we then interrogated whether polyQ-specific changes identified from omics profiling are relevant for neuronal toxicity in vivo in *Drosophila*.

Collectively, these analyses revealed a broad program associated with TBP expression/dosage that occurred independently of polyQ expansion and that was characterized by increased abundance of proteins associated with ribosomal function, translation, and mitochondrial processes, whereas tissue-specific polyQ-dependent responses included proteins involved in xenobiotic and redox pathways in the CNS and anabolic processes in skeletal muscle.

## RESULTS

### Validation of a *Drosophila* model of SCA17

TBP is a core transcription factor, but its pathogenic polyQ expansion in SCA17 likely rewires cellular programs beyond transcription via the capacity of aggregation-prone TBP-polyQ to affect the proteome via post-transcriptional mechanisms. TMT mass spectrometry offers an unbiased, quantitative readout of these downstream consequences at the level most proximal to the phenotype, i.e. the proteome, by capturing molecular changes that may be invisible or uncoupled from mRNA changes. Using tissue-resolved expression of human full-length TBP-WT and TBP-SCA17 in neurons (dissected heads; elav-Gal4) and skeletal muscle (dissected thoraces; Mef2-Gal4) in day-14 females, we profiled differential protein abundance vs. driver-only controls. This strategy resolves dosage vs. mutation-induced changes, identifying shared and divergent pathway vulnerabilities while identifying actionable targets.

At 14 days of age (week 2), adult female flies expressing TBP-WT or TBP-SCA17 did not show detectable locomotor phenotypes, as assessed by the RING assay (^19^; figure 1A summarizes our previously published findings; figure 1B shows mobility performance across 5 weeks). Their longevity was also indistinguishable from non-exogenous TBP-expressing controls at this stage (^19^; figure 1C), allowing us to dissect processes and pathways that precede symptom onset and are unlikely to be due to secondary events present after disease manifestation. We also quantified the effect of the expression of TBP-WT and TBP-SCA17 in fly eyes by using GFP-tagged CD8 as a correlate of retinal health and cell viability ^35–37^: these analyses indicated that expression of either form of human TBP reduced overall fluorescence (a proxy for cell viability), with TBP-SCA17 having a stronger impact (figure 1D). Collectively, these findings confirmed the validity of this model of SCA17.

### TMT mass spectrometry identifies the proteomic changes in the CNS and skeletal muscle of SCA17 models and controls

To quantify proteome-wide effects of TBP expression in vivo, we performed 18-plex TMT quantitative mass spectrometry on *Drosophila* lines expressing no exogenous TBP (control), human TBP-WT, or human TBP-SCA17 in neurons or in skeletal muscles (figure S1). We prepared biological triplicates consisting of 200 dissected heads (neuronal driver) per sample and 50 thoraces (muscle driver) per sample, respectively. This design leverages TMT multiplexing to measure relative protein abundance across all genotypes with high precision while controlling for developmental timing, genetic background, and driver effects. PCA analyses indeed indicated clustering of biological replicates, thus highlighting the validity of sample preparation and data collection (figures S1, S2, and S3).

### Venn diagrams and UpSet plots identify convergent and divergent changes induced by WT vs. mutant TBP in the CNS and skeletal muscle

To map convergence and divergence between normal TBP overexpression (WT) and disease-causing TBP (SCA17) relative to control (CTRL; driver only), we compared SCA17 vs. CTRL, WT vs. CTRL, and WT vs. SCA17 and summarized intersections as 3-set Venn diagrams for heads and thoraces (figure 1E). In heads, pairwise differences were dominated by WT vs. CTRL with 59 differentially expressed proteins (DEPs), suggesting strong TBP-dosage effects. Next, SCA17 vs. CTRL has 55 DEPs, and the WT vs. SCA17 comparison has 21 DEPs, providing clues into polyQ-dependent effects. The overlaps were dominated by the SCA17 vs. CTRL ∩ WT vs. CTRL intersection (34 proteins), with smaller pairwise-only regions involving WT vs. SCA17 (11 with SCA17 vs. CTRL and 6 with WT vs. CTRL) and a small triple intersection (2 proteins). Unique sets contributed 8 (SCA17 vs. CTRL), 17 (WT vs. CTRL), and 2 (WT vs. SCA17) proteins. In thoraces, pairwise DEP comparisons ranked as follows, highest to lowest: 92 DEPs in WT vs. CTRL, 41 in SCA17 vs. CTRL, and 20 in WT vs. SCA17, further underscoring gene dosage effects, but also highlighting polyQ-dependent outcomes. In terms of overlaps, the SCA17 vs. CTRL ∩ WT vs. CTRL intersection is comparable with the heads (28 proteins). Overlaps that include WT vs. SCA17 are modest (13 with WT vs. CTRL, 5 with SCA17 vs. CTRL), and the triple intersection is minimal (1 gene). Unique sets contributed 7 (SCA17 vs. CTRL), 50 (WT vs. CTRL), and 1 (WT vs. SCA17) protein(s).

We next quantified how the six differential-abundance sets intersect using an UpSet plot for head and thorax contrasts (WT vs. CTRL, SCA17 vs. CTRL, SCA17 vs. WT; figure 1F). In single sets, the largest were derived from WT vs. CTRL (with Thorax WT vs. CTRL the biggest and Head WT vs. CTRL next), followed by SCA17 vs. CTRL in each tissue; SCA17 vs. WT sets were smaller in both tissues. In terms of intersections, the largest were dominated by combinations that include CTRL-anchored contrasts (WT vs. CTRL and/or SCA17 vs. CTRL), often within the same tissue. Intersections that included WT vs. SCA17 were consistently smaller and many fine-grained combinations were single digit. Within-tissue intersections (e.g., Thorax WT vs. CTRL ∩ Thorax SCA17 vs. CTRL) exceeded cross-tissue intersections, indicating tissue-specific changes.

Based on these results, TBP-dosage biology (changes shared by WT vs. CTRL and SCA17 vs. CTRL) carries most of the signal, whereas SCA17-WT differences are limited. The UpSet view reinforces that these effects are strongly tissue-contextualized, with thorax and head contributing distinctive patterns rather than a single pan-tissue response or effect.

### Volcano plots highlight differentially abundant proteins in WT and mutant TBP vs. controls

Consistent with the intersection structures that we observed from Venn diagrams, volcano plots comparing WT to CTRL in heads showed a right-shift with many highly significant upregulated proteins linked to carbohydrate/lipid metabolism and biosynthesis (figure 2; e.g., Pepck2, PEPCK, tobi), together with stress/immune and signaling components (TepII, multiple gag proteins, CG10618, CG4962, CG3699, and CG10200). Several labels sat at −log_10_ P-value ≈ 8– 10 with log_2_FC ∼0.8–1.8, indicating a strong TBP-dosage dependency. Prominent decreases included secreted/innate and chemosensory factors (Bom23, BomBc2, AttC, DptA, and Obp99c) and small heat shock proteins (Hsp22), pointing to a shift from barrier/defense to metabolic output. The SCA17-to-CTRL comparison largely mirrored the WT pattern in heads: the same metabolic/stress machinery was elevated (Pepck2, PEPCK, TepII, and CG10618), with additional neural/xenobiotic signals (Rh5, Cyp6a23, and CG5106) among the positives; the secreted/chemosensory cohort (Bom23, BomBc2, Obp99c, and Lectin-galC1) was likewise reduced. Direct SCA17 vs. WT differences were fewer but specific: WT is higher for Cyp4p1, CG4408, CG2065, rdhB, Gmnt, CG16826, BomBc2, Obp83a/b, and CG7289, whereas spartin, were CG7829 are higher in SCA17. These differences suggest selective divergence in xenobiotic/retinoid handling, odorant-binding, and membrane-associated features superimposed on a shared metabolic backbone in neuronal tissue-rich heads.

**Figure 2:**
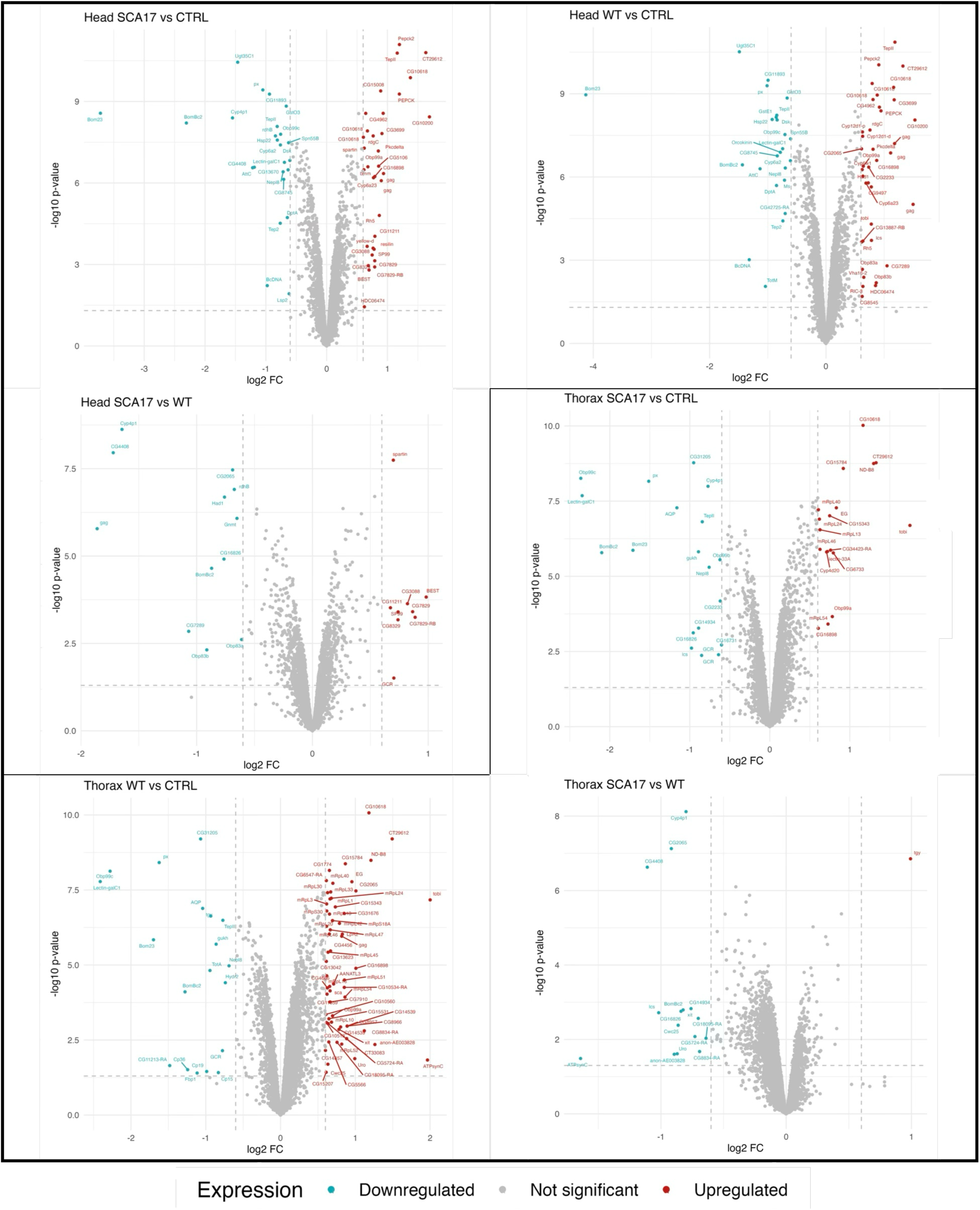
Volcano plots and pathway analyses of the proteomic changes. A) Volcano plots (P < 0.05 and |log_2_FC| ≥ 0.6) of heads (top) and thoraces (bottom) highlighting divergences across phenotypes.

In the muscle-enriched thoraces, the WT-to-CTRL comparison showed an even stronger anabolic/energetic signature: numerous mitochondrial ribosomal proteins (mRpL1/3/5/13/20/24/30–33, mRpS family), respiratory chain components (ND-B8), and anabolic enzymes (tobi, PEPCK) were markedly upregulated (many with −log_10_ P-value > 7 and log_2_FC ∼2). In parallel, secreted/chemosensory and innate proteins were reduced (Obp99c, Lectin-galC1, px, AQP, TepII, Bom23, BomBc2, and Nep1/8), consistent with prioritization of energy production and protein synthesis in flight muscle under TBP overexpression. SCA17 vs. CTRL in thoraces beared some resemblance with the WT profile: increased levels of mitochondrial ribosomal/respiratory components (mRpL40/46, CG15784, ND-B8) and decreased levels of the same barrier/defense cohort (Obp99c, Lectin-galC1, Bom23, BomBc2, AQP, px, gukh, TepII). However, the overall number of differences was markedly smaller than the WT-to-CTRL comparison. The SCA17 vs. WT comparison identified a compact set of differences with modest effect sizes (log_2_FC ∼0.4–1.3) but high significance for several features (−log10 P-value > 6): WT had higher levels than SCA17 of Cyp4p1, CG2065, CG4408, CG14934, Uro, CG5724-RA, CG8834-RA, and ATPsynC, while tgy was reduced in WT (it is higher in SCA17). These differences point to refined divergence in detoxification and nucleotide/lipid metabolism.

Collectively, the volcano plots reinforced the Venn/UpSet results, with TBP dosage dominating the proteomic response in both tissues. Relative to controls, TBP-WT and TBP-SCA17 exhibited shared increases in the abundance of proteins associated with metabolic and translational processes, alongside decreased abundance of proteins involved in secretory, innate immune, and chemosensory functions. This convergence was most striking in skeletal muscle, where mitochondrial ribosomal and respiratory proteins were higher while OBPs and lectins fell, consistent with energetic and biosynthetic prioritization in muscle. Neuronal tissues exhibited the same overall underlying framework with the additional modulation of neuronal/xenobiotic components. Moreover, the WT vs. SCA17 comparisons exposed focused, pathway-level differences. Thus, mutant TBP preserves the broad anabolic program associated with TBP overexpression, yet diverges in specific detoxification, retinoid/odorant-binding, and trafficking nodes, with differences that are more evident in heads vs. thoraces.

### GO and KEGG analyses identify enriched categories among modulated proteins

We next conducted Gene Ontology (GO) and KEGG analyses to place these proteomic shifts into pathway contexts (figure 3A). In heads, TBP dosage (WT vs. CTRL) was characterized at the biological-process level by enrichment for mitochondrial translation, mitochondrial gene expression, carboxylic acid, oxoacid, organic acid, and organophosphate metabolic processes, small-molecule biosynthesis, generation of precursor metabolites and energy, glutathione metabolism, and cuticle/chitin-associated categories, with additional representation of phototransduction and responses to oxygen-containing compounds (figure 3A). Molecular-function terms similarly emphasized structural constituents of the ribosome, heme, tetrapyrrole, and iron binding; oxidoreductase and monooxygenase activities; glutathione transferase activity; and structural constituents of cuticle and chitin-based cuticle (figure 3B). KEGG terms in head WT vs. CTRL included pentose and glucuronate interconversions, insect hormone biosynthesis, metabolism of xenobiotics by cytochrome P450, drug metabolism – cytochrome P450, drug metabolism – other enzymes, glutathione metabolism, cysteine and methionine metabolism, and arginine metabolism (figure 3C).

**Figure 3:**
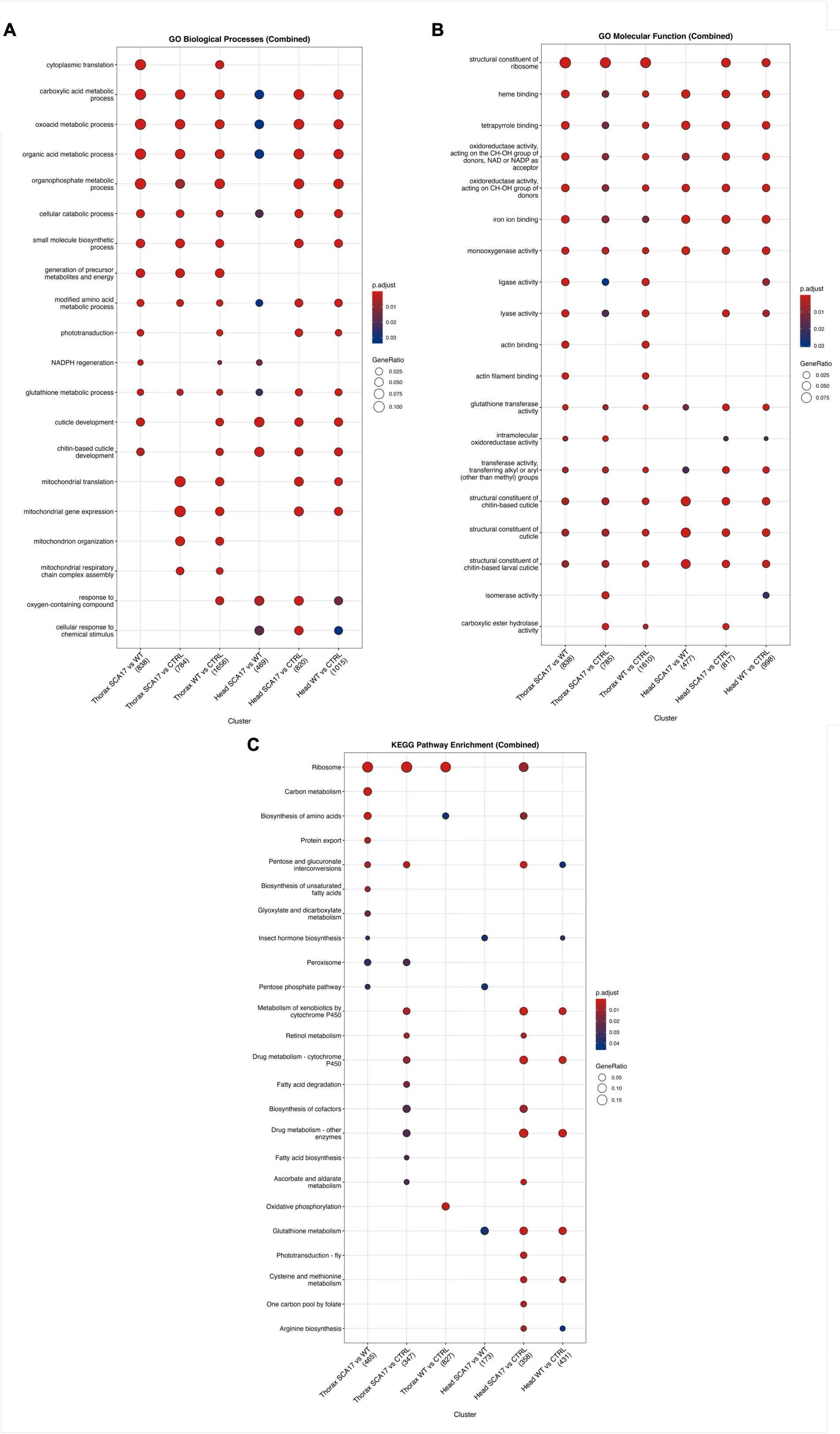
GO and KEGG analyses of the proteomic changes. A-C) GO Biological processes (A), Molecular functions (B) and KEGG analyses (C) of the information gathered by TMT proteomics.

SCA17 mutation relative to CTRL recapitulated the dosage program for biological processes (figure 3A). Molecular-function analysis also showed large overlaps, with terms again centered on ribosome structure, oxidoreductase and monooxygenase activities, heme/tetrapyrrole/iron binding, and glutathione transferase activity. One key difference was the emergence of carboxylic ester hydrolase activity in this comparison and the loss of isomerase activity compared to the WT-to-CTRL set (figure 3B). KEGG analysis retained several of the WT-associated pathways but broadened the xenobiotic-metabolic signature, highlighting ribosome, biosynthesis of amino acids, metabolism of xenobiotics by cytochrome P450, drug metabolism - cytochrome P450, drug metabolism - other enzymes, retinol metabolism, biosynthesis of cofactors, ascorbate and aldarate metabolism, phototransduction, glutathione metabolism, and cysteine and methionine metabolism (figure 3C). When isolating mutation effects from dosage (SCA17 vs. WT), biological processes no longer included induced mitochondrial translation and gene expression, and several metabolic processes, but instead introduced NADPH regeneration (figure 3A). Molecular functions retained oxidoreductase activities and structural components, but lost ribosome representation (figure 3B). KEGG highlighted three sole items: insect hormone biosynthesis, pentose phosphate pathway, and glutathione metabolism (figure 3C).

In thoraces, gene dosage (WT vs. CTRL) produced changes similar to those found in heads but with even broader biological process coverage, including cytoplasmic translation, mitochondrial gene expression and organization, mitochondrial respiratory chain assembly, and generation of precursor metabolites and energy (figure 3A). Molecular function analysis showed strong enrichment for structural constituents of the ribosome and translation factors, with pervasive oxidoreductase and monooxygenase functions, heme and iron binding, glutathione transferase activity, and structural constituents of cuticle and chitin (figure 3B). KEGG identified ribosome, biosynthesis of amino acids, and oxidative phosphorylation (figure 3C). The SCA17 mutation relative to CTRL retained some of these categories. Biological processes remained centered on mitochondrial translation and intermediary metabolism. However, enrichments for cytoplasmic translation, phototransduction, NADPH regeneration, and cuticle/chitin were absent (figure 3A). Molecular functions again featured ribosomal structure, oxidoreductase and monooxygenase activities, heme and iron binding, and glutathione transferase. Actin-related functions, translation factor activity, and proton channel activity were absent here (figure 3B). KEGG analysis preserved ribosome, and added pentose and glucuronate interconversions, cytochrome P450-related activities and drug metabolism, fatty acid degradation, retinol metabolism, and ascorbate and aldarate metabolism (figure 3C). Mutation relative to WT (SCA17 vs. WT) no longer enriched mitochondrial translation and gene expression, but reintroduced chitin/cuticle components (figure 3A). Molecular function largely resembled the WT-to-CTRL comparisons (figure 3B). KEGG enrichment for ribosome was retained, similar to what we observed with the other two comparisons (figure 3C). New additions in this analysis were carbon metabolism, protein export, biosynthesis of unsaturated fatty acids, insect hormone biosynthesis, and pentose phosphate pathway. Shared with the SCA17-to-CTRL comparison was the pentose and glucuronate interconversions and peroxisome, whereas biosynthesis of amino acids was shared with the WT-to-CTRL comparison (figure 3C).

Altogether, these analyses highlight TBP dosage as the dominant driver, impacting ribosome and mitochondrial translation, respiratory chain assembly, and broad carbon and amino acid metabolism, with broader effects in thoraces. SCA17 vs. CTRL largely displayed the same changes while adding KEGG categories for cytochrome P450, retinol, and xenobiotic pathways. Isolating mutation effects (SCA17 vs. WT) removed mitochondrial translation and ribosome categories in heads; heads shifted toward redox control with NADPH regeneration, pentose phosphate, insect hormone biosynthesis, and glutathione metabolism, whereas thoraces retained ribosome enrichment, reintroduce chitin and cuticle components, and expanded to carbon metabolism, protein export, and unsaturated fatty acid biosynthesis in SCA17 vs. WT. There were also tissue specificities in the responses seen in SCA17 vs. WT: heads showed phototransduction, thoraces emphasize fatty acid and central metabolism. We conclude that TBP dosage reprograms translation and bioenergetics in both tissues, while SCA17 superimposes selective redox and xenobiotic features that are more pronounced in heads than in thoraces.

### Changes in PTMs distinguish TBP WT vs. TBP SCA17 from controls

We used the JUMPptm approach^38–41^ to leverage the existing TMT dataset and extracted 22,096 modified peptides spanning acetylation, carboxylation, deamidation, dihydroxylation, mono-methylation, oxidation, phosphorylation, and ubiquitination. We first examined how significant PTM-carrying proteins (p < 0.05) distribute across the most prominent modification classes within each tissue (figure 4A). In heads, the signal was strongly partitioned by modification type and anchored by deamidation, which contained a large deamidation-only pool (752 proteins) and a major intersection with oxidation (96). Additional connectivity was comparatively limited and largely routed through dihydroxylation, including overlaps with deamidation (20), phosphorylation (12), and a small higher-order intersection (3). Phosphorylation and mono- methylation showed substantial class-specific components (for example, phosphorylation-only 36 and a phosphorylation–mono-methylation intersection of 5; 39 if deamidation is also included). Carboxylation in heads was entirely non-overlapping (70), similar to acetylation (95).

**Figure 4:**
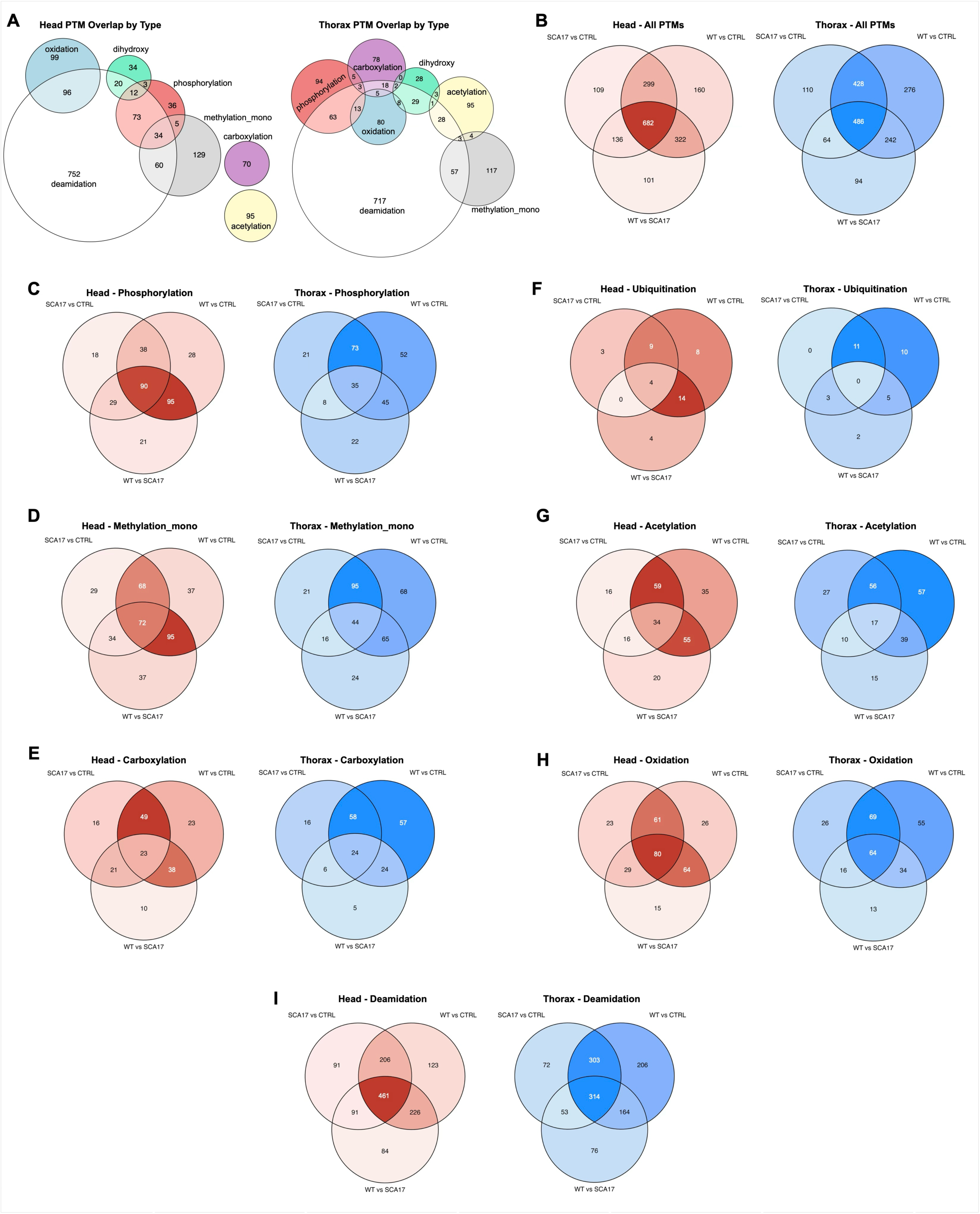
Post-translational modifications changes across genotypes and tissues. Euler plots (A) and Venn diagrams (B-I) of all the post-translational modifications that could be retrieved with JUMPptm from the TMT dataset, assessed by tissue and genotype.

In thoraces, deamidation again dominated with a large deamidation-only pool (717), but the Euler structure showed greater cross-class mixing than in heads (figure 4A). Phosphorylation intersected deamidation prominently (63), with additional higher-order overlap involving oxidation (13). Carboxylation and oxidation also participated in multi-class intersections (for example, carboxylation–oxidation, 5). Dihydroxylation (28) displayed additional overlap with acetylation (3). Acetylation retained a large stand-alone component (95) but also intersected both deamidation and mono-methylation, including an acetylation–deamidation overlap (28), an acetylation–mono-methylation overlap (4), and a three-way overlap among acetylation, mono-methylation, and deamidation (3). Mono-methylation likewise contained both a sizable stand-alone pool (117) and an overlap with deamidation (57). Together, these Euler summaries emphasize that, within each tissue, significant PTM-associated proteins distribute across partially overlapping modification classes rather than collapsing into a single shared set, with thoraces exhibiting more interconnection among major PTM types than heads (figure 4A).

We next quantified how PTM-associated proteins were shared across genotype comparisons within each tissue (figure 4B). In heads, WT vs. CTRL produced the largest set (1,463), with overlaps with SCA17 vs. CTRL (299) and WT vs. SCA17 (322). A large, shared core (682) was significant across all three head comparisons, alongside comparison-specific subsets (160 unique to WT vs. CTRL, 109 unique to SCA17 vs. CTRL, and 101 unique to WT vs. SCA17), indicating both broad commonality and comparison-distinct PTM structure in heads (figure 4B). Thoraces showed a similar organization with distinct magnitudes: WT vs. CTRL again yielded the largest set (1,432), with strong overlap with SCA17 vs. CTRL (428) and a substantial three- way shared core (486). Thoraces also displayed comparison-specific components (276 unique to WT vs. CTRL, 110 unique to SCA17 vs. CTRL, and 94 unique to WT vs. SCA17), consistent with a large common module alongside comparison-specific PTM shifts in this tissue (figure 4B).

Integrating the comparison-wise Venn diagrams for individual modification classes (figure 3C–I) showed that the extent of shared vs. comparison-specific signal varies by PTM type and tissue. In heads, phosphorylation was prominent across comparisons (WT vs. CTRL, 251; SCA17 vs. CTRL, 175; WT vs. SCA17, 235) and included a strong three-way shared core (90), with a large overlap between WT vs. CTRL, and WT vs. SCA17 (95) (figure 4C). Mono-methylation showed a similarly broad pattern (WT vs. CTRL, 272; SCA17 vs. CTRL, 203; WT vs. SCA17, 238), with 72 shared across all three and a large WT vs. CTRL–WT vs. SCA17 intersection (95) (figure 4D). Carboxylation in heads included 23 shared modified peptides across all three comparisons and strong pairwise overlaps (49 shared between WT vs. CTRL and SCA17 vs. CTRL, and 38 shared between WT vs. CTRL and WT vs. SCA17) (figure 4E). Ubiquitination was smaller in scale in heads (WT vs. CTRL, 35; SCA17 vs. CTRL, 16; WT vs. SCA17, 22) but retained a detectable three-way core (4) (figure 4F). Acetylation in heads was robust (WT vs. CTRL, 183; SCA17 vs. CTRL, 125; WT vs. SCA17, 125), with 34 shared across all three and substantial overlap between WT vs. CTRL and WT vs. SCA17 (55) (figure 4G). Oxidation and deamidation also contributed large, comparison-spanning signals in heads, each with sizable three-way shared cores (80 for oxidation; 461 for deamidation) (figure 4H–I).

In thoraces, phosphorylation remained prominent (WT vs. CTRL, 205; SCA17 vs. CTRL, 137; WT vs. SCA17, 110) but with a smaller three-way shared core (35) than in heads (figure 4C). Mono-methylation again showed broad involvement (WT vs. CTRL, 272; SCA17 vs. CTRL, 176; WT vs. SCA17, 149), with 44 shared across all comparisons (figure 4D). Carboxylation in thoraces included 24 modified peptides that are shared across all three comparisons, and strong pairwise overlap between WT vs. CTRL and SCA17 vs. CTRL (58) (figure 4E). Ubiquitination remained limited (WT vs. CTRL, 26; SCA17 vs. CTRL, 14; WT vs. SCA17, 10) and lacked a three-way shared core (0), indicating more comparison-specific behavior in thoraces (figure 4F). Acetylation was again robust (WT vs. CTRL, 169; SCA17 vs. CTRL, 110; WT vs. SCA17, 81), with a smaller shared core than in heads (17) (figure 4G). Oxidation and deamidation remained major contributors in thorax, with substantial three-way shared cores (64 for oxidation; 314 for deamidation) (figure 4H–I).

Overall, these PTM-focused analyses reveal broad remodeling in both tissues, with large, shared cores across comparisons when PTMs are considered in aggregate, and with phosphorylation, mono-methylation, acetylation, oxidation, and deamidation each contributing substantial comparison-spanning components. The Euler summaries further indicate that, within each tissue, significant modified proteins distribute across partially overlapping modification classes, with thoraces exhibiting more interconnection among major PTM types than heads (figure 4A–I).

Consistent with the extensive overlap patterns across modifications (figure 4), volcano plots in figure 5 revealed broad bidirectional changes in PTMs across tissues and genotype comparisons. In heads, WT vs. CTRL produced significant changes spanning a wide range of effect sizes, with a visibly prominent enrichment of upregulated features. The heads SCA17 vs. CTRL comparison similarly displayed both increased and decreased modified peptides and an apparent bias toward positive fold changes. In contrast, the direct heads SCA17 vs. WT comparison was more constrained, with fewer significant features overall and a tighter clustering around the center, consistent with a smaller residual set of mutation-associated post-translational modification differences when WT gene dosage is held constant.

**Figure 5:**
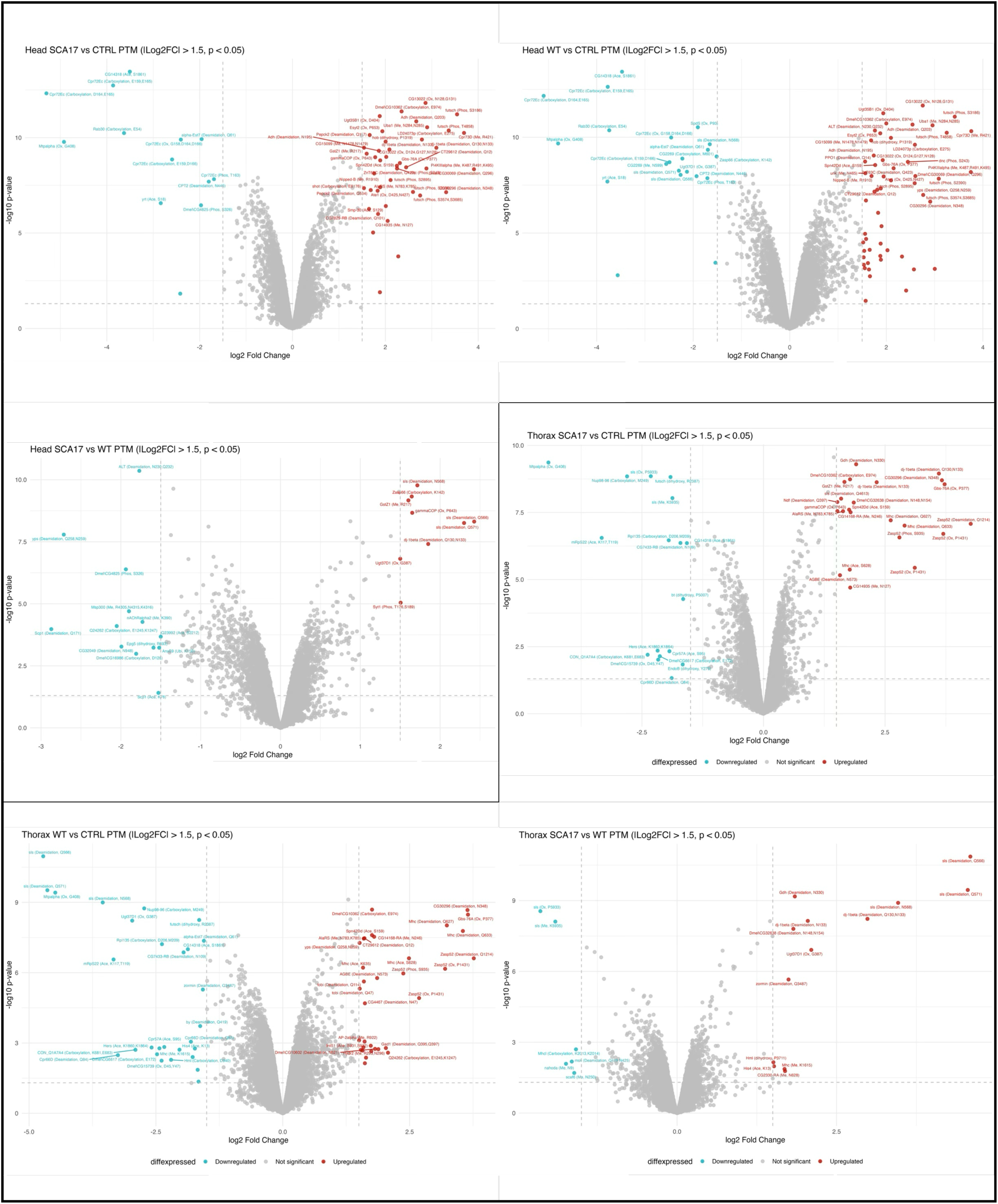
Volcano plots of post-translational modifications changes across genotypes and tissues. Post-translational modification data from heads (top) and thoraces (bottom) plotted as volcano plots with |log_2_FC| > 1.5 and P < 0.05.

Thoraces showed the same overall hierarchy. WT vs. CTRL again displayed modified peptides distributed broadly in both directions. Thoracic SCA17 vs. CTRL also displayed an extensive pattern of significant changes, with multiple features exceeding the fold-change thresholds on both sides of the plot. However, thoracic SCA17 vs. WT was the most restricted of the thorax comparisons, with comparatively few significant peptides and a more centralized distribution despite the persistence of both up- and downregulated PTMs. Collectively, these volcano plots reinforce that most PTM changes are observed in comparison with CTRL (WT vs. CTRL, and SCA17 vs. CTRL), whereas the direct mutation contrast (SCA17 vs. WT) captures a smaller, more selective subset of changes in both tissues.

### Orthogonal RNA-seq analyses reveal transcriptional changes caused by WT and mutant TBP vs. controls

In parallel with TMT proteomics, we conducted RNA-seq analyses using the exact same tissues. We examined pairwise and three-way comparisons for head and thoracic RNA-seq data (SCA17 vs. CTRL, WT vs. CTRL, and WT vs. SCA17; figure 6; PCA analyses are in figure S4). Across tissues, Venn analyses of RNA-seq data revealed broader remodeling in thoraces vs. heads. In heads, SCA17 vs. CTRL, WT vs. CTRL, and WT vs. SCA17 yielded 621, 987, and 632 differentially expressed genes (DEGs), respectively, with modest pairwise overlaps (SCA17 vs. CTRL ∩ WT vs. CTRL: 263; SCA17 vs. CTRL ∩ WT vs. SCA17: 109; WT vs. CTRL ∩ WT vs. SCA17: 270) and a small core shared by all three contrasts (33; figure 5A). In thoraces, DEG changes were far larger (3068 in SCA17 vs. CTRL; 1360 in WT vs. CTRL; and 2563 in WT vs. SCA17). We observed an extensive convergence between SCA17 vs. CTRL and WT vs. SCA17 (1286), substantial overlap with WT vs. CTRL (717 and 156 in the other pairwise intersections), and a central shared set of 287 genes. We also observed unique segments in both tissues (e.g., head: 216/421/220; thorax: 778/200/834 for the three contrasts; figure 6A), indicating genotype-specific programmatic rewiring. Overall, head changes were more distributed and distinct across comparisons, whereas thoracic changes show pronounced, shared perturbations between TBP overexpression and the SCA17 mutation. These results are consistent with stronger dosage-driven and mutation-convergent effects in muscle-rich tissue.

**Figure 6:**
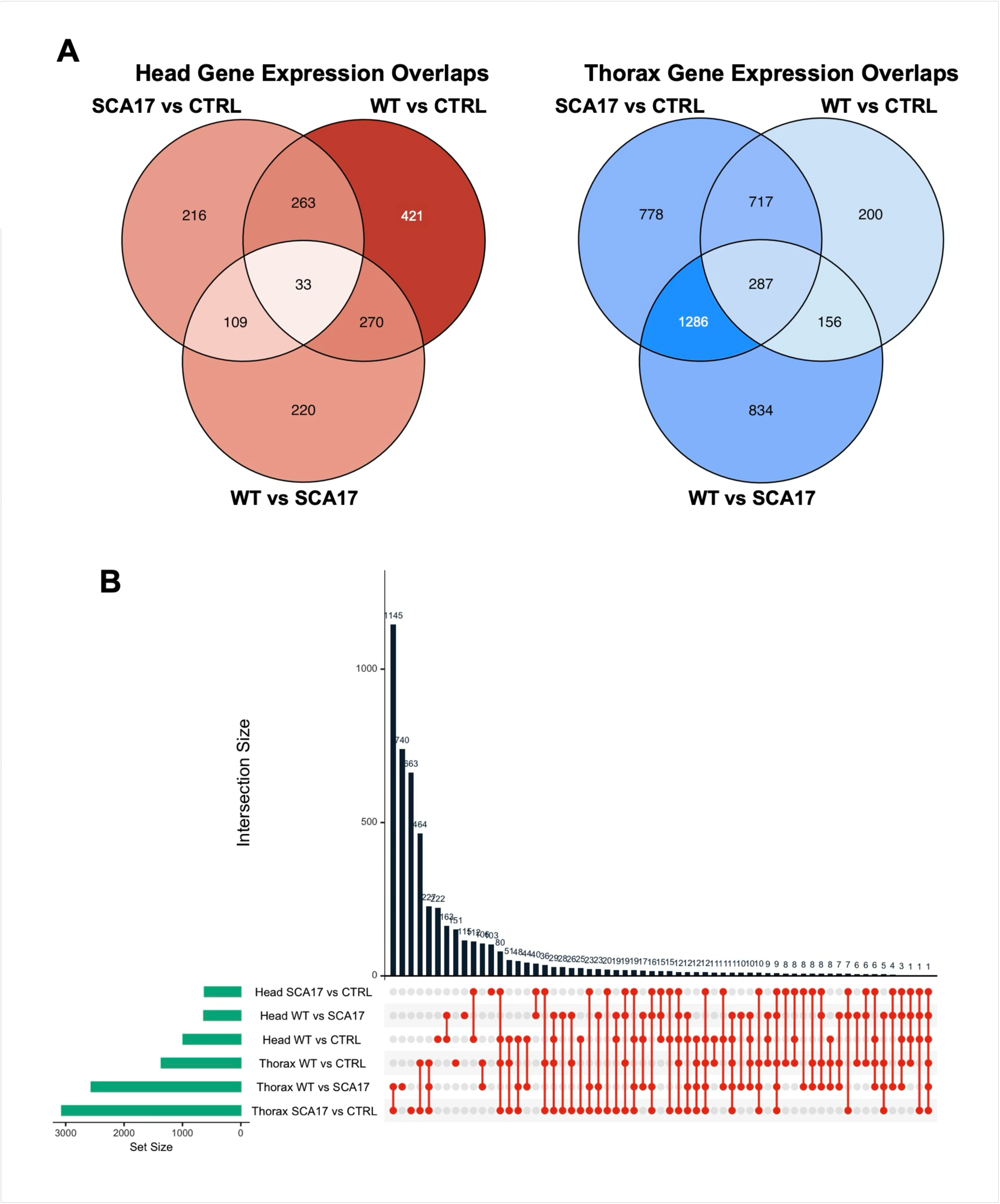
RNA-seq result convergences and divergences across samples and tissues. A) Venn diagrams summarizing transcript similarities and differences among genotypes and tissues. B) UpSet plot quantifying the transcript intersections across the six genotypes and both tissues.

We next applied an UpSet analysis to enumerate and prioritize shared vs. unique cohorts within and across tissues for downstream analyses (figure 6B). The UpSet analysis across six contrasts showed thorax-dominated overlaps. The largest intersection (n=1145) was shared by thorax SCA17 vs. CTRL, and thorax WT vs. SCA17, followed by thorax-unique sets for SCA17 vs. CTRL (n=740), WT vs. SCA17 (n=663), and SCA17 vs. CTRL and WT vs. CTRL thoracic groups (n=464; figure 6B). Subsequent intersections were markedly smaller (∼222, 163, 151, 120, 106, 103, 80, then ≤51) and included various cross-tissue combinations. By comparison to thorax-based outcomes, head-only and head/thorax overlapped are modest (<200), with minimal elements spanning many contrasts (figure 6B). Overall, thoraces harbored the dominant shared remodeling, while head changes were fragmented.

To visualize effect size and direction underlying these intersections, we constructed volcano plots for each contrast in heads and thoraces (figure 7). Volcano plots reinforced tissue-specific burdens. Thoraces showed more marked differences in the SCA17 vs. CTRL and SCA17 vs. WT comparisons, suggesting extensive induction and repression consistent with mutation-convergent and dosage-linked remodeling. Thorax WT vs. CTRL showed a smaller but significant set, aligning with TBP-dosage effects. In heads, contrasts yielded comparatively fewer significant species and narrower fold-change ranges. Prominent outliers were concentrated in thoracic contrasts, whereas head contained sparser extreme responders. Together, these distributions highlight thorax as the primary locus of strong, shared transcriptomic remodeling, with head changes being more modest and dispersed.

**Figure 7:**
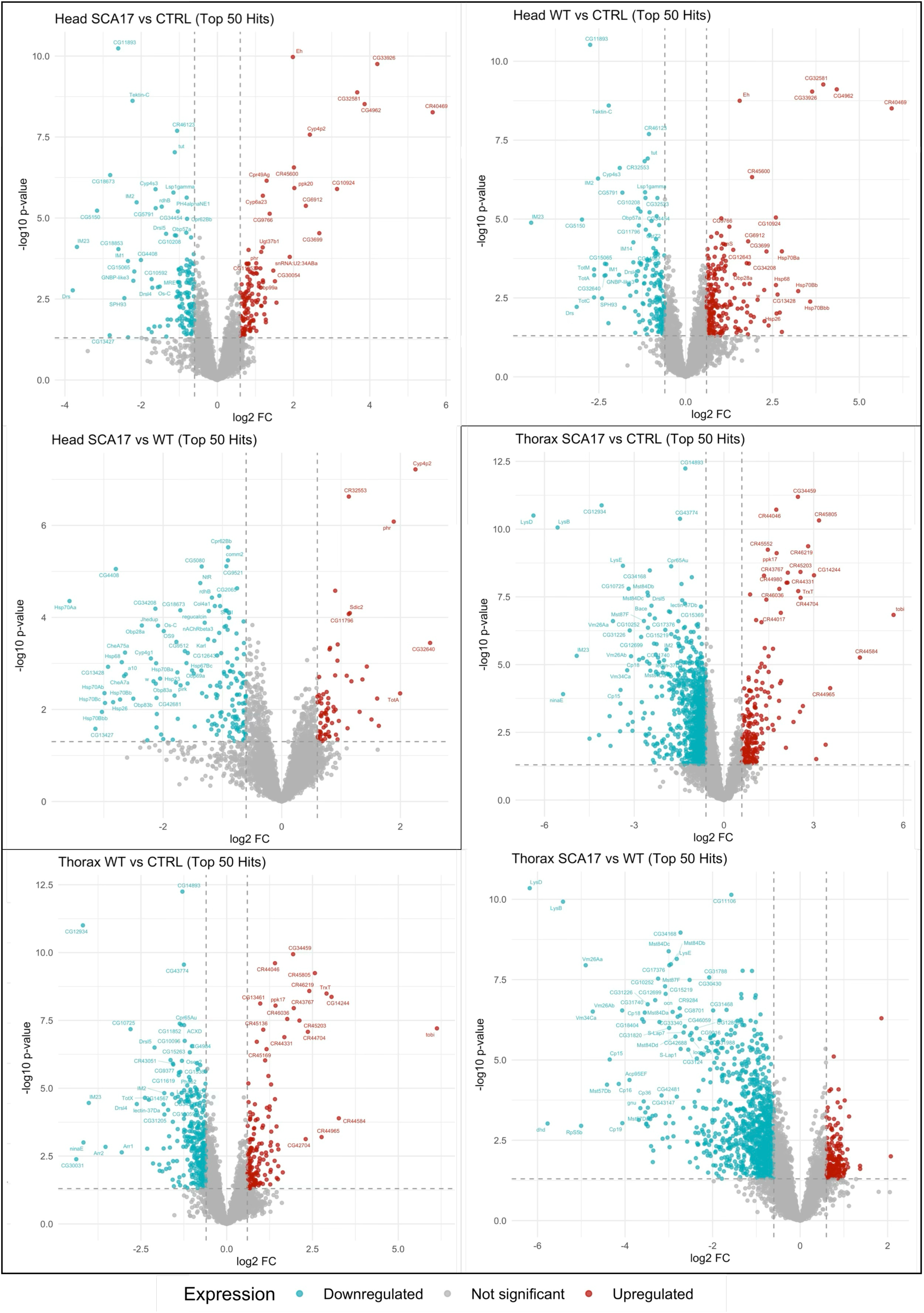
**Volcano plots of RNA-seq result across samples and tissues.**

We next focused on GO (figure 8A, B) and KEGG (figure 8C) analyses. Across head-derived datasets, DEGs were enriched in processes linked to sensory signaling, cell cycle regulation, immune defense, and proteostasis. GO molecular function analysis supported these trends by identifying enrichment for protein folding processes (e.g., ATP-dependent protein folding chaperone, heat shock protein binding), signaling (odorant binding), and pyrophosphatase activity (figure 8B). KEGG pathways (figure 8C) identified immune response elements (Toll and IMD signaling pathways), proteostasis (protein processing in ER), cell signaling (endocytosis), and cell cycle regulation (spliceosome), mirroring the GO terms for Biological Processes (figure 8A).

**Figure 8:**
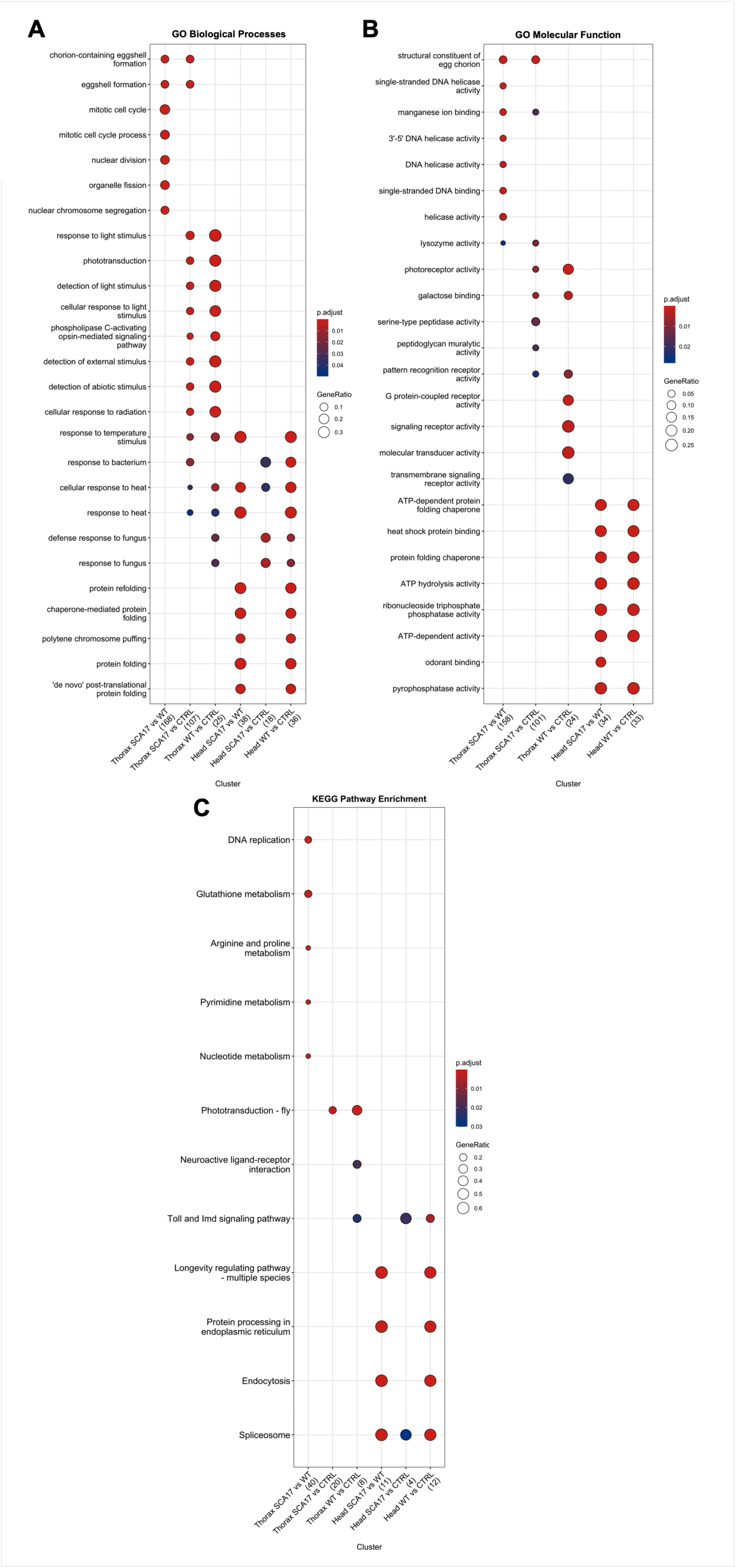
GO and KEGG analyses of the transcriptomic changes. A-C) GO Biological processes (A), GO Molecular functions (B) and KEGG analyses (C) of the information gathered by RNA-seq. We did not observe significant differences in the SCA17-to-CTRL comparison for ‘Molecular Function.’

As we focused on pairwise comparisons, differences were more pronounced in WT-to-CTRL and SCA17-to-CTRL sets, indicative of TBP gene dosage effects, than in the SCA17-to-WT comparison in Biological Processes (figure 8A). Intriguingly, the SCA17-to-CTRL comparison yielded no significant differences in Molecular Function analysis. The other two comparisons were nearly identical, with odorant binding being the sole difference: present in the SCA17 vs. WT comparison, but not in WT vs. CTRL, indicating that this function is potentially dependent on polyQ length in fly heads. KEGG analyses showed only two differences in the SCA17-to-CTRL comparison: Toll and Imd signaling pathways and spliceosome. Both these pathways were present in the WT-to-CTRL comparison, but the Toll and Imd signaling pathway category was absent in the SCA17-to-WT comparison. Lastly, several pathways were shared in their enrichment when comparing SCA17-to-WT with WT-to-CTRL sets. Together, the head transcriptomes indicate coordinated remodeling of sensory transduction, innate immunity, chaperone networks, and cell cycle programs that vary in their specific enrichments in a gene dosage- and polyQ-dependent manner.

Thorax-based outcomes were more varied overall, perhaps because of the larger number of DEGs retrieved from this tissue. In thoraces, distinct effects that are gene-dosage-dependent vs. those that are polyQ-length dependent emerged in Biological Processes (figure 8A). Cellular responses to foreign agents and various stimuli were especially TBP dose-dependent, whereas processes such as cell cycle were especially polyQ length-dependent. These distinctions were not fully mirrored by Molecular Function analysis (figure 8B). While there were functions that were enriched in the gene dosage comparisons (WT-to-CTRL and SCA17-to-CTRL), such as photoreceptor activity, galactose binding and pattern recognition receptor activity, other functions were distinctly enriched in only one set or another (WT-to-CTRL: G protein-coupled receptor, signaling receptor activity, molecular transducer activity, and transmembrane signaling receptor activity; SCA17-to-CTRL: peptidoglycan muralytic activity, serine-type protease activity, and lysozyme activity). The SCA17-to-WT comparison yielded several cell cycle and transcription-related activities, as well as lysozyme activity. Structural constituents of egg chorion also emerged as enriched in SCA17-to-WT and SCA17-to-CTRL comparisons. KEGG analyses yielded clear distinctions in each pairwise comparison (figure 8C). In WT-to-CTRL comparisons, Toll and Imd signaling pathway, neuroactive ligand-receptor interaction, and phototransduction were enriched, with phototransduction being the sole component shared with the SCA17-to-CTRL comparison, highlighting this process as especially TBP dose-dependent. The SCA17-to-WT comparison led to the enrichment of several pathways that appear to be polyQ-dependent, including DNA replication and cell metabolism. Collectively, the thorax-based data underscore similarities with processes and pathways in heads (responses to stimuli, albeit of different kinds, and immune responses), as well as differences, such as cell cycle-related events and reproductive processes.

Based on these collective results across tissues, TBP gene dosage appears to be the primary driver of transcriptional changes, with thoraces showing larger and more convergent changes than heads. Venn and UpSet analyses highlight thorax-dominated overlaps, including a large common core between SCA17 vs. CTRL and WT vs. SCA17, consistent with mutation effects aligning to a dosage-driven program in muscle-rich tissue. Overall, gene dosage establishes the dominant and shared transcriptomic framework, particularly in the thorax, while polyQ length imposes selective overlays that are more evident in the thorax vs. the head.

Comparing outcomes from both omics approaches indicates that TBP dosage is the primary driver and that the thorax displays the largest, most convergent changes. RNA-seq Venn, UpSet, and volcano plots show extensive shared remodeling in thoraces, especially for SCA17 vs. CTRL and for WT vs. SCA17, whereas heads show smaller, more fragmented changes.

TMT data agree on this hierarchy and directionality: WT and SCA17 relative to CTRL share a strong anabolic program with higher ribosomal and mitochondrial components and lower secreted, innate, and chemosensory proteins, most strikingly in the thorax. In both approaches, SCA17 vs. CTRL broadly overlaps with WT vs. CTRL changes, and SCA17 vs. WT yields fewer but specific differences, indicating that polyQ effects are largely superimposed on a dosage-established framework.

The omics diverge in emphasis, consistent with regulatory vs. effector layers. TMT results accentuate bioenergetic and translational outputs in the thoracic flight muscle and pinpoint polyQ-dependent shifts in detoxification, retinoid, odorant-binding, and trafficking modules that are more evident in heads. RNA-seq data from the same tissues highlight remodeling of sensory signaling, innate immunity, proteostasis, and cell-cycle programs, with clearer polyQ-length signatures in the thorax, including DNA replication and metabolic pathways, and a head-specific molecular function category for odorant binding in SCA17 vs. WT. Together, the datasets strongly suggest that dosage reprograms translation and mitochondrial metabolism in both tissues, that the thorax shows the strongest shared response, and that polyQ length adds selective, tissue-tuned overlays that are transcriptionally most apparent in the thorax and post-transcriptionally (i.e., at the proteome level) most apparent in the head.

### Direct RNA-to-protein comparisons reveal proteomic changes arising from transcriptional vs. post-transcriptional mechanisms

To directly quantify the relationship between transcriptional and proteomic changes, we plotted mRNA log_2_ fold changes against protein log_2_ fold changes for all gene products detected by both platforms (∼4,500 per tissue) and computed Spearman rank correlations for each pairwise comparison (figure 9, supplemental data S2). Points were classified by significance category: significant in both RNA and protein (“Both sig”), RNA-only significant, PROT-only significant, or not significant in either (NS).

**Figure 9:**
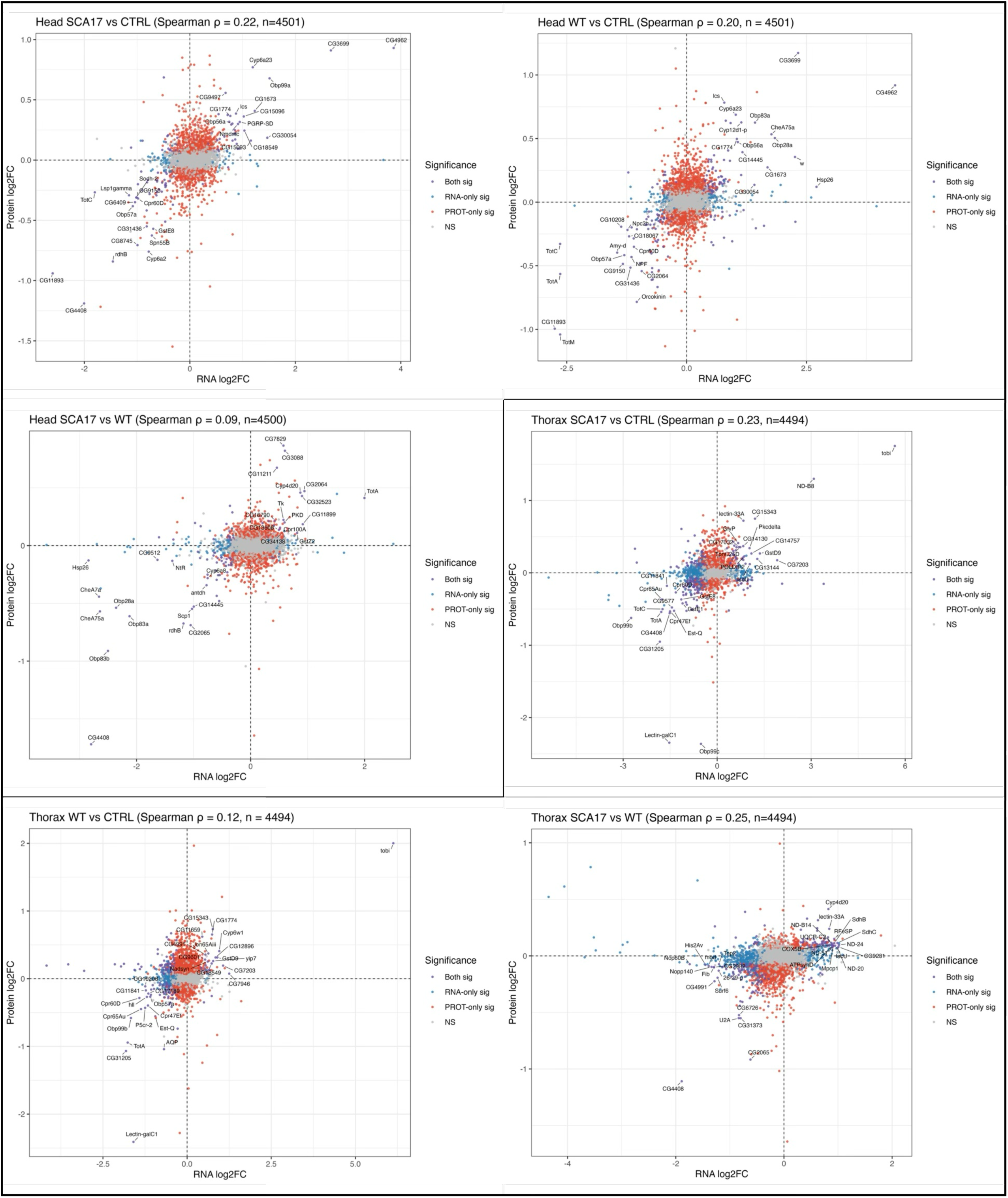
RNA-to-protein correlation across genotypes and tissues. Scatter plots of matched mRNA log_2_FC (x-axis) vs. protein log_2_FC (y-axis) for each pairwise comparison in heads and thoraces. Points are color-coded by significance category: Both sig (significant in both platforms), RNA-only sig, PROT-only sig, or NS (not significant in either). Spearman rho and the number of matched gene products are indicated in each panel title.

In CTRL-anchored comparisons, mRNA and protein fold changes showed modest positive correlations in both tissues: head SCA17 vs. CTRL (rho = 0.22; figure 9), head WT vs. CTRL (rho = 0.20), thorax SCA17 vs. CTRL (rho = 0.23), and thorax WT vs. CTRL (rho = 0.12). These positive correlations indicate that, when either form of TBP is compared to driver-only controls, directional changes in transcript abundance are partially preserved at the protein level, and that many proteomic changes do not arise from corresponding transcriptional changes. The correlation was stronger for SCA17 vs. CTRL in both tissues (0.22 in heads, 0.23 in thoraces), suggesting that the combined dosage-plus-mutation signal produces more concordant RNA-protein changes than dosage alone. Thorax WT vs. CTRL yielded the weakest positive correlation (0.12), suggesting that a substantial fraction of the TBP-dosage program in muscle is post-transcriptionally buffered or amplified asymmetrically at the protein level.

The direct SCA17 vs. WT comparisons also produced positive Spearman correlations: head SCA17 vs. WT (rho = 0.09; figure 9) and thorax SCA17 vs. WT (rho = 0.25). These results indicate that mutation-dependent differences between TBP-WT and TBP-SCA17 retain a positive, although tissue-dependent, correspondence between mRNA and protein levels. This relationship is weak in heads (rho = 0.09), consistent with relatively limited cross-platform concordance in the direct comparison, but stronger in thoraces (rho = 0.25), where the direct SCA17-WT contrast shows the strongest correlation observed across all six comparisons.

Across all six comparisons, “Both sig” features (significant in both RNA and protein) were a minority relative to platform-specific significant features, indicating that much of the biologically relevant signal captured by each platform is not necessarily recapitulated by the other. RNA-only significant genes were abundant across all comparisons, particularly in the thorax, consistent with the larger DEG counts observed by RNA-seq in this tissue. Protein-only significant features were more evenly distributed across comparisons. Together, these correlation analyses provide a quantitative framework for the cross-platform observations described above: TBP dosage effects are modestly concordant between RNA and protein, whereas the mutation-specific divergence between WT and SCA17 is also positively correlated across platforms, with especially strong concordance in the muscle-enriched thorax vs. the head.

### Validation of omics hits identifies evolutionary-conserved modifiers of SCA17 pathology

We next tested whether selected mutation-associated proteomic changes were functionally relevant for SCA17 pathology. We examined four candidate modifiers in flies expressing TBP-SCA17 pan-neuronally. We selected them primarily from changes that distinguished TBP-SCA17 from TBP-WT, thereby focusing the validation on the narrower polyQ-dependent component of the proteomic response rather than on the broader effects of TBP dosage. Among the four candidates, spartin and CG2065 were prioritized because they changed in opposite directions in the proteomic dataset: spartin was increased in TBP-SCA17 relative to TBP-WT, whereas CG2065 was decreased. We therefore tested whether reducing spartin would ameliorate the phenotype and whether further reduction of CG2065 would exacerbate it. We also included Cyp4d20 and Tsp42Ea, for which the proteomic results did not suggest a comparably clear functional direction, to assess the specificity of these predictions. We assessed locomotor performance in flies expressing TBP-SCA17 in all fly neurons in the presence or absence of these four candidates.

As shown in figure 10A, RNAi targeting spartin supported the prediction derived from its increased abundance in the SCA17 proteome. Although spartin RNAi had little detectable effect at weeks 1 and 4, it significantly improved climbing performance by week 5 relative to flies expressing TBP-SCA17 alone. This delayed benefit suggests that increased spartin contributes to the progressive neuronal dysfunction caused by polyQ-expanded TBP rather than representing a uniformly protective response. CG2065 RNAi (which targets the NADP retinol dehydrogenase), however, did not produce the anticipated worsening. Instead, reducing CG2065 significantly improved climbing, with the strongest benefit evident at week 4 and a smaller but persistent improvement at week 5. Thus, the reduced abundance of CG2065 in the SCA17 proteome may represent an adaptive protective response. Neither Cyp4d20 RNAi nor a loss of function mutation of Tsp42Ea produced a significant change to TBP-SCA17 control in this assay. Together, these data provide functional support for spartin and CG2065 as modifiers of neuronal TBP-SCA17 toxicity, while also illustrating that the direction of an omics change does not necessarily reveal whether it is a pathogenic or compensatory protective response. Spartin and CG2065 may influence SCA17-associated neuronal dysfunction via their previously reported functions in protein trafficking, membrane homeostasis, and proteostasis.

**Figure 10:**
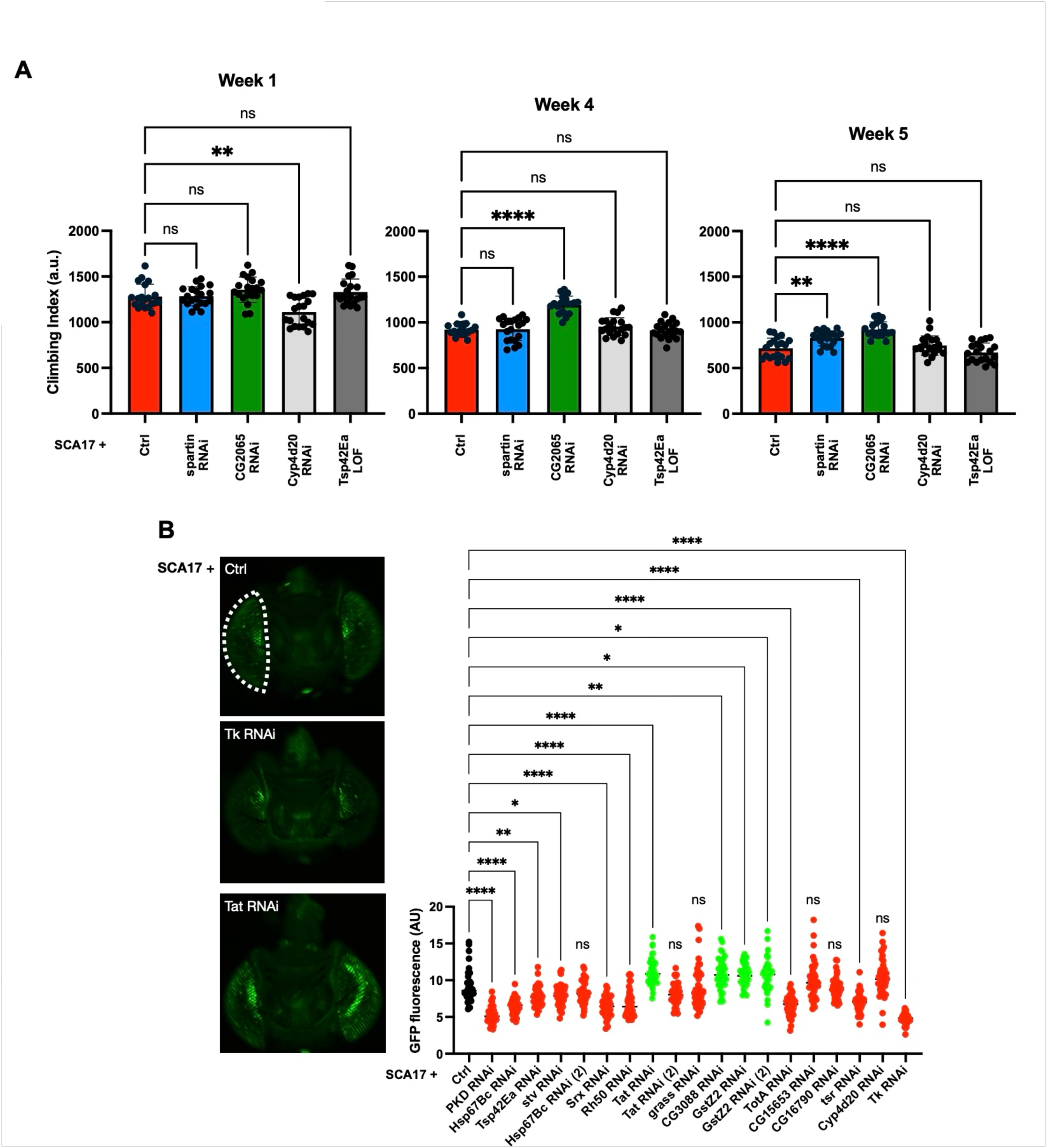
Validation of candidate genes identifies modifiers of SCA17 pathology. A) Motility assessment of adult flies expressing the noted transgenes in all fly neurons at weeks 1, 4, and 5. N=100 per cohort. Statistics: Brown-Forsythe and Welch’s ANOVA tests. *P<0.05; **P<0.01; ***P<0.001; ****P<0.0001. B) Effect of reducing the noted candidate genes on GFP fluorescence in eyes co-expressing TBP-SCA17 and CD8-GFP. Left: representative images are shown for TBP-SCA17 alone, TBP-SCA17 with Tk RNAi and TBP-SCA17 with Tat RNAi. Right: Statistics: one-way ANOVA with Dunnett’s multiple comparisons test vs. TBP-SCA17 alone. N=20 per genotype. *P<0.05; **P<0.01; ****P<0.0001; ns, not significant. Red color indicates reduced GFP fluorescence compared to the TBP-SCA17 alone; green color indicates increased fluorescence.

Lastly, we further validated these hits by using a CD8-GFP eye reporter (figure 1D). In this assay, GFP fluorescence (indicative of cell viability) is reduced by TBP-WT and this occurs more strongly with TBP-SCA17. Because this readout is quantitative and scalable, it allowed us to interrogate more candidates than the motility assay. We drew candidates from the head (neuron-enriched) SCA17-vs.-WT comparison, prioritized by cross-platform congruence (table S2). We selected two classes: proteins changed significantly and in the same direction at both the transcript and the protein level (PKD, Tsp42Ea, Srx, Tat, CG3088, GstZ2, TotA, CG15653, CG16790, Cyp4d20, and Tk), and a discordant class in which protein abundance increased while the corresponding mRNA levels declined (Hsp67Bc, stv, Rh50, grass, and tsr), a pattern which implicates post-transcriptional regulation. Functionally, this set includes regulators of redox, glutathione, xenobiotic and stress handling (Srx, GstZ2, Cyp4d20, and TotA); tyrosine metabolism (Tat); chaperone-dependent proteostasis (Hsp67Bc, and stv); cytoskeletal and membrane/trafficking components (tsr, PKD, and Tsp42Ea); and neuronal or secreted signaling factors (Tk, Rh50, and grass). Every candidate taken from the congruence analysis was more abundant in TBP-SCA17 than in TBP-WT heads.

As summarized in figure 10B, RNAi-dependent targeting of individual candidates in flies co-expressing TBP-SCA17 significantly altered eye GFP fluorescence for 13 of the 19 RNAi lines tested (12 of the 16 genes; some were targeted using two independent RNAi lines). Three genes behaved as suppressors, increasing fluorescence relative to TBP-SCA17 alone: Tat, CG3088, and GstZ2, for which two independent RNAi lines gave concordant results. Nine genes behaved as enhancers, decreasing fluorescence: PKD, Hsp67Bc, Tsp42Ea, stv, Srx, Rh50, TotA, tsr, and Tk. Four genes produced no significant change: grass, CG16790, CG15653, and Cyp4d20. For interventions with 2 distinct RNAi lines, results were concordant for GstZ2, directionally concordant but not statistically different for Hsp67Bc, and divergent for Tat RNAi lines.

Our original expectation for the targets that we tested with CD8-GFP was different. Since the levels of each of the targets was higher in TBP-SCA17 vs. -WT in congruent omics data, we would have predicted that knockdown of each of them would have been more likely to increase fluorescence. Only three out of the sixteen tested did so. The five discordant ones (mRNA down/protein up: Hsp67Bc, stv, Rh50, grass, and tsr) behaved uniformly opposite to prediction (i.e., knockdown worsened the phenotype) which suggests that post-transcriptional upregulation of these proteins in SCA17 is protective.

Taken together with the locomotor data in figure 7A, where reducing spartin or CG2065 improved climbing in flies expressing TBP-SCA17 pan-neuronally, the CD8-GFP-derived results provide in vivo functional validation for a substantial portion of the polyQ-dependent omics signature, since 12 of 16 candidates and 4 of 4 candidates tested across the two assays measurably modified TBP-SCA17 toxicity. These results also indicate that the direction of an omics-derived change does not predict its functional contribution: increased abundance in TBP-SCA17 corresponds to pathogenic drivers in some cases (spartin, Tat, CG3088, GstZ2) but to protective, largely post-transcriptional responses in others (Hsp67Bc, stv, Srx, Tk, PKD, tsr, and Rh50).

## DISCUSSION

The multi-omic findings presented here build directly on the phenotypic characterization of these same *Drosophila* SCA17 lines that we reported previously^19^. Earlier, we observed that expression of either wild-type TBP or SCA17-linked polyQ-expanded TBP in specific tissues or ubiquitously was problematic, with the pathogenic version causing greater toxicity. We also noted that TBP-SCA17 accumulated progressively higher levels of SDS-resistant, aggregation-prone species relative to TBP-WT, and that the disease-associated protein preferentially localized to the nucleus^19^. These observations are consistent with established features of polyQ disease pathology^14^. In this study, we extended these phenotypic observations to the molecular level through tissue-resolved TMT proteomics, PTM profiling, and RNA-seq, identifying molecular pathways that are reshaped by TBP-SCA17 in the early phases of disease etiology, i.e. before the onset of phenotypic differences in longevity and motility assays.

By profiling day-14 adults at a pre-symptomatic stage, our proteomic analyses revealed that TBP dosage itself, independent of polyQ expansion, is a dominant driver of proteomic remodeling in both neuronal and muscle-enriched tissues. Based on Venn diagrams and UpSet analyses in both heads and thoraces, the WT vs. CTRL comparison produced the largest number of DEPs, followed by SCA17 vs. CTRL, with SCA17 vs. WT yielding the fewest changes. The overlaps between WT vs. CTRL and SCA17 vs. CTRL dominated the intersection structure, while within-tissue intersections exceeded cross-tissue intersections, something consistent with strongly tissue-contextualized effects rather than a single pan-tissue response. For example, both TBP-WT and TBP-SCA17 exhibited increased abundance of proteins associated with carbohydrate and lipid metabolism, stress responses, signaling, translation, and mitochondrial function, together with reduced abundance of proteins involved in innate immunity and chemosensory processes.

This dosage-driven program could not be mechanistically clarified by prior phenotypic analyses, in which TBP-WT and TBP-SCA17 effects were consistently aligned in direction, with the SCA17 version generally leading to greater severity^19^. The present, tissue-resolved analyses resolve this relationship: a shared TBP-dosage backbone operates in both conditions, upon which a mutation-specific divergence is superimposed. The SCA17 vs. WT comparison exposed focused divergencies, including those in xenobiotic and retinoid handling, odorant-binding proteins, membrane and trafficking modules that are observed in heads, and refined differences in detoxification and nucleotide and lipid metabolism that are found in thoraces. GO and KEGG analyses reinforced and extended these findings at the pathway level. In both tissues, TBP dosage drove enrichment for mitochondrial translation, mitochondrial gene expression, respiratory chain assembly, and broad carbon and amino acid metabolism, with a broader scope in thoraces than in heads. SCA17 vs. CTRL largely overlaid the dosage program while adding KEGG signals for cytochrome P450, retinol metabolism, and xenobiotic pathways.

When SCA17 was compared to TBP-WT to distinguish mutation associated changes from the shared TBP response, mitochondrial translation and the ribosome were no longer enriched jn heads; instead the head response shifted toward redox associated pathways, including NADPH regeneration, the pentose phosphate pathway, insect hormone biosynthesis, and glutathione metabolism. The SCA17-to-WT comparison in thoraces retained ribosome enrichment and expanded to carbon metabolism, protein export, biosynthesis of unsaturated fatty acids, and showed enrichment for chitin and cuticle components. Together, these data indicate that SCA17 does not impose a wholly new pathway architecture; rather, it reshapes the TBP-dosage program in a tissue-dependent manner, narrowing the head response toward redox/xenobiotic handling and reducing the full anabolic-bioenergetic engagement achieved by WT in the thorax.

The PTM analyses added a further dimension to these findings, revealing broad post-translational remodeling that extended beyond changes in protein abundance. In both tissues, deamidation anchored the PTM landscape, contributing the largest stand-alone pools and the most extensive cross-class connectivity. However, deamidation and oxidation can occur during sample preparation and may not exclusively reflect biological processes. In particular, non-enzymatic deamidation can occur under the alkaline conditions used during protein digestion for mass spectrometry analyses, requiring further downstream validation. Phosphorylation, mono-methylation, acetylation, and oxidation each contributed substantial comparison-spanning components, with large three-way shared cores across all genotype comparisons, particularly in heads. Thoraces exhibited more interconnection among major PTM types than heads, a finding that parallels the broader scope of proteomic and transcriptomic remodeling observed in muscle-enriched tissue. The genotype-comparison hierarchy at the PTM level mirrored the protein-abundance data: contrasts against CTRL (WT vs. CTRL and SCA17 vs. CTRL) produced the most expansive PTM remodeling, whereas SCA17 vs. WT captured a smaller, more selective subset. Both tissues retained comparison-specific PTM subsets alongside their shared cores, indicating that TBP dosage and polyQ expansion each contribute distinct post-translational signatures. The prevalence of deamidation, a modification associated with protein aging and turnover, alongside phosphorylation and mono-methylation, is consistent with the metabolic and proteostatic reprogramming identified at the protein-abundance level; it may reflect accelerated protein turnover, altered signaling cascades, and metabolic stress imposed by TBP overexpression.

Orthogonal RNA-seq analyses from the same tissues provided a complementary regulatory perspective. In contrast to the proteomic data, where head and thorax DEP numbers were more comparable, transcriptomic remodeling was far more extensive in thoraces, with DEG counts reaching over 3,000 in the SCA17 vs. CTRL compared to several hundred DEGs in heads.

Venn and UpSet analyses highlighted thorax-dominated overlaps, with the largest intersection shared between thorax SCA17 vs. CTRL and thorax SCA17 vs. WT, consistent with mutation effects converging on a dosage-driven program in skeletal muscle. Head transcriptomic changes were smaller and more fragmented across comparisons. GO and KEGG analyses of the RNA-seq data resolved tissue-specific features that complemented the proteomic findings.

The integration of proteomic and transcriptomic datasets revealed informative platform-specific emphases consistent with the distinction between regulatory and effector molecular layers. TMT proteomics accentuated bioenergetic and translational outputs, particularly in thorax flight muscle, and pinpointed mutation-specific shifts in detoxification, retinoid, odorant-binding, and trafficking modules that were more evident in heads. RNA-seq data highlighted remodeling of sensory signaling, innate immunity, proteostasis, and cell cycle programs, with clearer polyQ-length signatures in thorax including DNA replication and metabolic pathways. Direct RNA-to-protein correlation analyses provided quantitative support for these observations: CTRL-anchored comparisons yielded modest positive Spearman correlations, confirming partial concordance between transcript and protein changes driven by TBP dosage or combined dosage-plus-mutation effects. However, the direct SCA17 vs. WT comparison produced positive correlations in both tissues: a weak correlation in heads and the strongest correlation among the six panels in thoraces. The stronger thoracic concordance indicates that mutation-dependent RNA and protein changes are more closely aligned in muscle-enriched tissue than in heads, despite the broader transcriptomic remodeling observed in thoraces. This observation underscores the value of orthogonal, multi-omic profiling: neither platform alone captures the full spectrum of TBP dosage and mutation effects, and the tissue-specific post-transcriptional filtering of polyQ-dependent signals represents an additional layer of biology that warrants further investigation.

For our functional validation tests, we focused on spartin and CG2065 because their abundance differed between TBP-SCA17 and TBP-WT in heads and because they represent distinct biological processes. Spartin was increased in TBP-SCA17 relative to TBP-WT, whereas CG2065 was reduced. We predicted that reducing spartin would be beneficial and that further reducing CG2065 would worsen neuronal dysfunction. Indeed, pan-neuronal spartin RNAi improved locomotor performance, supporting our prediction that increased spartin contributed to progressive TBP-SCA17 toxicity. CG2065 RNAi also improved locomotion; in this case, the direction of its proteomic change did not directionally predict the effect of experimental suppression.

Spartin is an evolutionarily conserved organelle-associated protein found at endosomes, lipid droplets, mitochondria, and neuronal membranes^42–48^. It participates in endosomal trafficking^49,50^, synaptic regulation^48,51^, lipid-droplet turnover, and mitochondrial Ca^2+^ homeostasis. It is also required presynaptically for normal neurotransmission at the neuromuscular junction, and it can restrain synaptic growth through endocytic regulation of BMP (bone morphogenic protein) signaling^43^. These functions place spartin at an intersection of membrane trafficking, synaptic homeostasis, lipid metabolism, and mitochondrial regulation, all processes that suggest a possible role in the neuronal resilience to an aggregation-prone nuclear protein such as TBP-SCA17. However, the improvement that we observed with spartin RNAi suggests that its elevation becomes maladaptive as TBP-SCA17-induced toxicity progresses. One possibility is that excess spartin shifts endosomal or receptor trafficking in a manner that compromises synaptic maintenance. Alternatively, changes in lipid-droplet dynamics or mitochondrial Ca^2+^ regulation could alter the ability of neurons to accommodate for chronic proteostatic and metabolic stress.

The other disease modifier, CG2065, provides a mechanistic link to the retinoid and redox pathways highlighted by the head proteome. CG2065 encodes an ER membrane-associated, NADP-dependent all-trans-retinol dehydrogenase of the short-chain dehydrogenase/reductase family. It catalyzes the oxidation of all-trans-retinol to all-trans-retinal while converting NADP+ to NADPH and has experimentally supported roles in retinal metabolism in multiple species^52–55^. Its reduced abundance in TBP-SCA17 relative to TBP-WT initially suggested that loss of this activity might be detrimental. Instead, further reduction improved locomotion, leading us to hypothesize that the proteomic decrease represents a compensatory, protective response.

Lower CG2065 activity could limit production of retinal or alter local retinoid flux, thereby reducing a toxic metabolic or signaling consequence of TBP-SCA17 expression. Because the reaction catalyzed by CG2065 also affects the NADP+/NADPH balance, CG2065 reduction could influence redox homeostasis in a setting where NADPH regeneration, pentose phosphate levels, and glutathione metabolism were already selectively implicated.

The spartin and CG2065 results indicate two different relationships between omics changes and disease mechanisms. Increased spartin abundance was directionally informative: reducing it ameliorated late locomotor dysfunction. Reduced CG2065 abundance was also functionally informative, but potentially as part of an adaptive response that could be enhanced experimentally. These outcomes emphasize that an observed molecular change may be pathogenic, protective, or accessory and hence not related to disease progression.

The results that we collected from the CD8-GFP screen diverged from what the omics alone would have predicted. Of the candidates that had increased levels in TBP-SCA17 relative to TBP-WT, some behaved as though their upregulation was pathogenic, whereas others behaved as though it was protective, since reducing their levels increased cytotoxicity. Regarding the induction of an adaptive response, Hsp67Bc and stv encode the *Drosophila* orthologs of HSPB8 and BAG3, partners in the chaperone-assisted selective autophagy module; also, Srx (sulfiredoxin) reverses peroxiredoxin hyperoxidation, all of which are activities expected to buffer proteotoxic and oxidative stress. Tissue provenance was likewise informative. Cyp4d20 and Tsp42Ea were the only candidates with concordant transcript and protein increases in both neuronal and muscle-enriched tissues, and they were two of the least effective modifiers here, producing no significant change (Cyp4d20) or a modest enhancement (Tsp42Ea); some of the strongest modifiers (Tk, Pkd, Srx, and Hsp67Bc) displayed head-restricted changes with no corresponding thoracic changes.

Prior investigations into SCA17 provide important context for interpreting these multi-omic results. In transgenic SCA17 mice, polyQ expansion in TBP enhanced its interaction with the general transcription factor TFIIB while reducing TBP dimerization^5^. This altered interaction coincided with reduced TFIIB occupancy at the Hspb1 promoter and suppression of the small heat shock protein HSPB1, a factor critical for neuronal and axonal integrity. Overexpression of HSPB1, and to a lesser extent TFIIB, alleviated mutant TBP-induced defects, placing the TBP– TFIIB–HSPB1 axis among the earliest defined molecular lesions in SCA17^56^. The pre-symptomatic head proteome described here is consistent with this framework: the SCA17 shifts toward CYP–glutathione redox and retinoid pathways, together with the loss of mitochondrial translation enrichment in the SCA17 vs. WT comparison, may reflect downstream consequences of selective transcriptional dysregulation rather than a global transcriptional failure. The head RNA-seq data reinforce this interpretation, showing enrichment for chaperone networks and ER protein processing that may represent compensatory responses to the proteostatic burden imposed by misfolded TBP.

Additional work indicated that polyQ expansion reduces the intrinsic affinity of TBP for TATA box DNA and that TBP fragments lacking an intact C-terminal DNA-binding domain accumulate in SCA17 mouse brain ^6^. These fragments formed nuclear aggregates and induced neurotoxicity despite impaired promoter engagement, supporting the idea that mutant TBP can disrupt transcriptional regulation through altered protein-protein interactions and complex assembly, independent of canonical DNA binding. In this context, the dosage-driven proteomic program described here, including robust upregulation of translation and mitochondrial oxidative phosphorylation even in TBP-WT, indicates that increased TBP abundance alone is sufficient to remodel cellular metabolism. The additional SCA17-specific divergences likely arise from aberrant interactions introduced by the expanded polyQ tract.

The thorax-derived data further align with prior observations of muscle pathology in SCA17. In a knock-in mouse model, large polyQ expansions in TBP caused primary skeletal muscle degeneration characterized by disrupted myofibril architecture, centralized nuclei, reduced fiber cross-sectional area, and mitochondrial abnormalities^17^. Mechanistically, polyQ expansion reduced the association of TBP with the muscle-specific transcription factor MyoD, impairing MyoD-dependent transcriptional programs and reducing expression of muscle structural genes^17^. Our *Drosophila* thorax data provide a pre-degenerative complement to these findings at several molecular levels. At the proteomic level, TBP-SCA17 already exhibits a lower anabolic set point relative to TBP-WT, with WT surpassing SCA17 in ribosome, oxidative phosphorylation, and amino acid metabolism pathways. At the transcriptomic level, thoraces showed the most extensive remodeling of any tissue–platform combination, with polyQ-dependent enrichment of cell cycle and DNA replication pathways that were not observed in heads. The convergence of attenuated anabolic proteomics with expanded cell-cycle transcriptomics in SCA17 thoraces suggests that muscle-intrinsic molecular deficits are detectable at a pre-symptomatic stage, and that the skeletal muscle compartment may be particularly sensitive to the combined effects of TBP dosage and polyQ expansion.

The neuronal proteomic and transcriptomic signatures identified here also connect to therapeutic studies implicating ER stress in SCA17 pathogenesis^16^. In previous work, mutant TBP induced ER stress and reduced expression of the neurotrophic factor MANF in the cortex and cerebellum, while piperine treatment restored MANF levels, reduced ER stress-associated toxicity, and improved neuronal survival. The head RNA-seq enrichment for protein processing in the ER and HSP binding is consistent with engagement of this axis in our pre-symptomatic assessment. The CG2065 result adds a further potential connection between the ER membrane, retinoid metabolism, and NADP(H)-linked redox regulation, whereas spartin links the phenotype to endosomal, synaptic, lipid-droplet, and mitochondrial biology. These candidates therefore extend the mechanistic framework beyond transcriptional dysregulation alone.

Earlier work supports strategies aimed at lowering mutant TBP levels or reversing its transcriptional consequences^18^. The targets identified across our datasets align with these principles and extend them through tissue-resolved, multi-layered evidence. Modulation of protein translation and mitochondrial quality control addresses the dosage-driven anabolic load evident in both proteomic and transcriptomic data. Regulation of CYP–glutathione detoxification pathways and retinoid metabolism address the redox and signaling alterations observed predominantly in neuronal tissue across both platforms. The polyQ-dependent cell cycle and DNA replication signatures in the thorax transcriptomes identify a previously unrecognized axis that may contribute to muscle vulnerability. Finally, the broad PTM remodeling, with its tissue-specific patterns of deamidation, phosphorylation, and mono-methylation, suggests that post-translational regulatory networks represent an additional layer of actionable mechanistic and potentially therapeutic biology in SCA17.

Taken together, the convergence of quantitative proteomics, PTM profiling, and RNA-seq with prior mechanistic studies provides a coherent and testable framework linking TBP dysfunction to tissue-specific vulnerability. By separating TBP dosage effects from polyQ-dependent perturbations across three molecular layers, our findings reveal that dosage reprograms translation and bioenergetics in both tissues, that polyQ expansion adds selective, tissue-tuned overlays that manifest themselves differently at the protein vs. transcript levels, and that PTMs are extensively remodeled in a tissue- and genotype-dependent manner. Collectively, this multi-omic framework offers a foundation for developing targeted, tissue-aware therapeutic strategies in SCA17.

## METHODS

### *Drosophila* genetics and husbandry

Transgenic lines expressing full-length human TBP were used: UAS-TBP-WT (Q25) and UAS-TBP-SCA17 (Q63), each HA-tagged and integrated at attP2, as described previously (single-copy, phiC31-mediated insertion)^19^. Pan-neuronal and muscle expression were driven with elav-Gal4 and Mef2-Gal4, respectively. Genetic controls were generated by crossing w^1118^ to the corresponding Gal4 drivers (driver-only controls). Flies were maintained on standard food (2% agar, 10% sucrose, 10% yeast) at 25°C, 12:12 h light:dark, and ∼40% relative humidity. Crosses were set at controlled density; adult progeny were synchronized within 12 h of eclosion. Unless stated otherwise, only females were used and assayed at day 14 of adulthood (chosen to coincide with robust Gal4-UAS expression and to avoid early post-eclosion physiological shifts), following the approaches described in our prior work^19,57,58^. The lines used for validation experiments are: elav-Gal4 (BDSC#458), Mef2-Gal4 (BDSC#27390), GMR-Gal4 (BDSC#8605), w^11^^18^ (BDSC#3605), UAS-mCD8::GFP (BDSC#5130), UAS-TBP-WT (Todi Lab generated), UAS-TBP-SCA17 (Todi Lab generated), UAS-spartin RNAi (BDSC#37499), UAS-CG2065 RNAi (BDSC#55283), UAS-Cyp4d20 RNAi (BDSC#77341), UAS-Tsp42Ea LOF (BDSC#13628), UAS-TspEa RNAi (BDSC#39044), UAS-Pkd RNAi (BDSC#28717), UAS-Hsp67Bc RNAi (BDSC#35452, BDSC#42607), UAS-stv RNAi (BDSC#42564), UAS-Srx RNAi (BDSC#43662), UAS-Rh50 RNAi (BDSC#50666), UAS-Tat RNAi (BDSC#76065, BDSC#51470), UAS-grass RNAi (BDSC#51920), UAS-CG3088 RNAi (BDSC#55145), UAS-GstZ2 RNAi (BDSC#68076, BDSC#55634), UAS-TotA RNAi (BDSC#58257), UAS-CG15653 RNAi (BDSC#62435), UAS-CG16790 RNAi (BDSC#64493), UAS-tsr RNAi (BDSC#65055), and UAS-Tk RNAi (VDRC#KK103662).

### Survival analysis and negative geotaxis

Cohorts of age-matched females (grouped in vials of ∼20) were maintained under the conditions described above and transferred to fresh food every 2–3 days. Death events were scored at each transfer until all flies died, and survival curves were generated for each genotype/driver combination as in Patel et al., 2023. Survival was analyzed by log-rank (Mantel–Cox) test with multiplicity-adjusted pairwise comparisons vs. driver-only controls. Locomotion was quantified by the Rapid Iterative Negative Geotaxis (RING) assay. Briefly, sets of five vials (20 females/vial) were tapped to the bottom, and the climbing height after 2 s was recorded. Twenty images were captured per week/group, and all images were batch-processed in ImageJ to limit processing variation. Group means were compared by linear models on trial-averaged heights; when longitudinal data were collected, slopes and intercepts were compared by linear regression as in^19^.

### CD8-GFP assessment

CD8-GFP-based experiments were conducted as previously described^35–37^. Briefly, GMR-Gal4 was used to express membrane-targeted UAS-CD8-GFP in the fly retina together with UAS-TBP-WT or UAS-TBP-SCA17. For the modifier screen, flies carrying GMR-Gal4, UAS-CD8-GFP, and UAS-TBP-SCA17 were crossed to the indicated RNAi lines and respective controls. Adult female progenies were aged for 14 days, at which time their heads were dissected, mounted on microscope slides, and imaged with a 4X objective on an Olympus BX51 fluorescence microscope, using identical acquisition settings. The ImageJ freehand selection tool was used to outline retinal areas and record the mean GFP fluorescence.

### Mass spectrometry

For sample preparation and TMT labeling for each genotype/driver and tissue (heads, thoraces), day-14 female samples were collected on ice and homogenized with zirconium beads in 8 M urea with 50 mM HEPES pH 8.5. After addition of 0.5% sodium deoxycholate, debris was pelleted. Protein concentration was determined by short Coomassie gels vs. BSA standards. Aliquots (100 µg) were digested with LysC (1:100, 2 h) then diluted to 2 M urea and digested with trypsin (1:50, ≥3 h). Reduction/alkylation steps were performed (DTT 1 mM, 30 min; IAA 10 mM, 30 min, dark; quench with 30 mM DTT). Peptides were acidified (1% TFA), desalted on C18 cartridges, dried, and labeled for TMTpro 18-plex (Thermo Fisher Scientific A52045) following our optimized protocol^59–63^.

For fractionation and LC–MS/MS, equal amounts of labeled peptides were pooled, desalted, and fractionated by basic-pH reverse-phase HPLC (pH 8.0, XBridge C18, 4.6 mm × 25 cm, 3.5 µm). Fractions were dried, reconstituted in 5% formic acid, and analyzed by nano-LC–MS/MS (Ultimate 3000 + Q-Exactive HF). Data-dependent acquisition used Orbitrap MS1/MS2 scans at 60k resolution with standard AGC targets, isolation windows, HCD NCE 32, and 10 s dynamic exclusion^58^.

Raw files were processed with JUMP (tag-based hybrid search) against the UniProt *Drosophila* reference proteome plus reversed decoys^40,64^. Search parameters: 25 ppm precursor/product tolerances, fully tryptic with ≤2 missed cleavages; static TMT on Lys and peptide N-termini (+304.20715 Da); Met oxidation as variable (+15.99492 Da). PSMs were filtered to ∼1% protein-level FDR. Protein abundance was computed by summing reporter-ion intensities across matched PSMs.

JUMPptm analyses to recover information on peptides modified by post-translational modifications (PTMs) from TMT mass spectrometry data was done as previously reported^41,65–67^. However, deamidation of asparagine and glutamine residues and oxidation of methionine residues can occur during protein extraction, digestion, and sample handling. Therefore, identification of these modifications by mass spectrometry does not necessarily establish their biological origin.

Principal component analyses were performed on normalized reporter intensities (R prcomp). Differential abundance was computed as log_2_ fold-change (contrast of interest) with two-sided testing; proteins meeting P < 0.05 and |log2FC| ≥ 0.6 were called differentially abundant (Nuga et al., 2024). Set overlaps were visualized with GeneOverlap (genome size background appropriate for *Drosophila*; odds ratios and P-values reported). Over-representation analyses used clusterProfiler with Gene Ontology (Biological Process and Molecular Function) and KEGG, Benjamini–Hochberg adjusted FDR < 0.05. Figures (volcano plots, heatmaps) were generated in R (ggplot2, ComplexHeatmap). All statistical tests are specified with each assay above. Data met assumptions of the tests used or were analyzed with nonparametric alternatives. Multiple-testing correction used Benjamini–Hochberg when applicable. Biological replicate definitions and n values are provided in the figure legends; independent crosses/biological samples were used for each replicate. The TMT mass spectrometry data have been deposited to the ProteomeXchange Consortium via the PRIDE partner repository.

### RNA sequencing

RNA sequencing was done and analyzed as previously described^36,57,68–71^. In brief, RNA-seq samples were prepared using TRIzol (Ambion #15596018), followed by RNA extraction by isopropanol precipitation from the aqueous phase. Libraries were constructed from 1 µg of total RNA using the Illumina TruSeq RNA Sample Prep v2 Kit, following the manufacturer’s instructions. Sequencing was carried out on the Illumina NovaSeq 6000 platform, generating 100-bp paired-end reads. These reads were quality trimmed and filtered to retain sequences with a Phred score of 20 or higher and lengths of at least 50 base pairs. The processed reads were then aligned to the *Drosophila melanogaster* reference genome (BDGP6/dm6) using CLC Genomics Workbench version 20.0.4 (Qiagen). Gene expression levels were quantified as transcripts per million (TPM) using the CLC RNA-Seq Analysis tool. Differential expression was evaluated through non-parametric ANOVA, employing Kruskal-Wallis and Dunn’s multiple comparisons tests on log-transformed TPM data within Partek Genomics Suite version 7.0 (Partek Inc.). The RNA-seq datasets generated in this study are available in the NCBI Gene Expression Omnibus (GEO) GEO (GSE339281).

### Statistics

Statistics were completed and data were visualized using GraphPad Prism version 11. Detailed statistical analyses are provided in the relevant Methods sections and within the figure legends, where applicable.

## DATA AVAILABILITY

All relevant data are included in the figures, tables and supplemental files. Omics data are available as indicated in the relevant Methods sections. Any data can be requested by contacting the corresponding authors and will be made readily available.

## ACKNOWLEDGMENTS

We thank Dr. Oluwademilade Nuga for her help with initial RNA-Seq analyses. This study was partially supported by NIH grant R01NS086778 (SVT).

## AUTHOR CONTRIBUTIONS

MVP: Conceptualization, methodology, software, validation, formal analysis, investigation, data curation, writing – original draft, writing – review and editing, visualization.

KR: Conceptualization, methodology, software, validation, formal analysis, investigation, data curation, writing – original draft, writing – review and editing, visualization.

FHH: Investigation, validation, writing – review and editing, visualization TK: Investigation, validation, visualization

AS: Investigation, validation

YDW: Methodology, software, formal analysis, data curation WW: Methodology, software, formal analysis, data curation

AAH: Methodology, software, formal analysis, data curation, writing – review and editing SP: Methodology, software, formal analysis, data curation, supervision, writing – review and editing

FD: Conceptualization, methodology, formal analysis, resources, writing – original draft, writing – review and editing, supervision, project administration.

SVT: Conceptualization, methodology, formal analysis, resources, writing – original draft, writing – review and editing, supervision, project administration, funding acquisition.

---

**Supplemental data S1: List of the proteins captured by Venn diagrams in figure 1E**.

**Supplemental data S2: List of congruence outcomes.**

**Figure S1:**
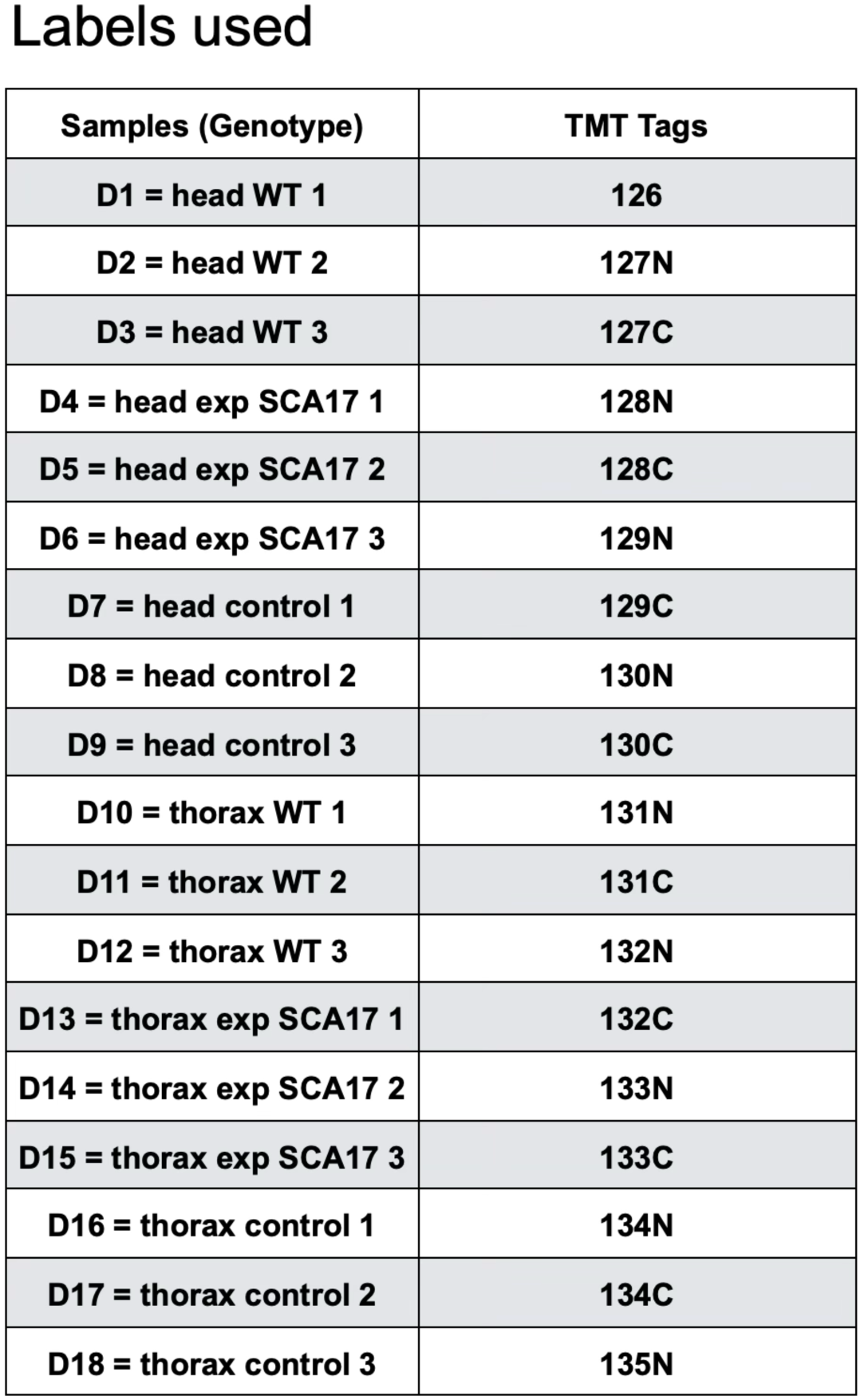
Proteomic TMT tags used for each sample.

**Figure S2:**
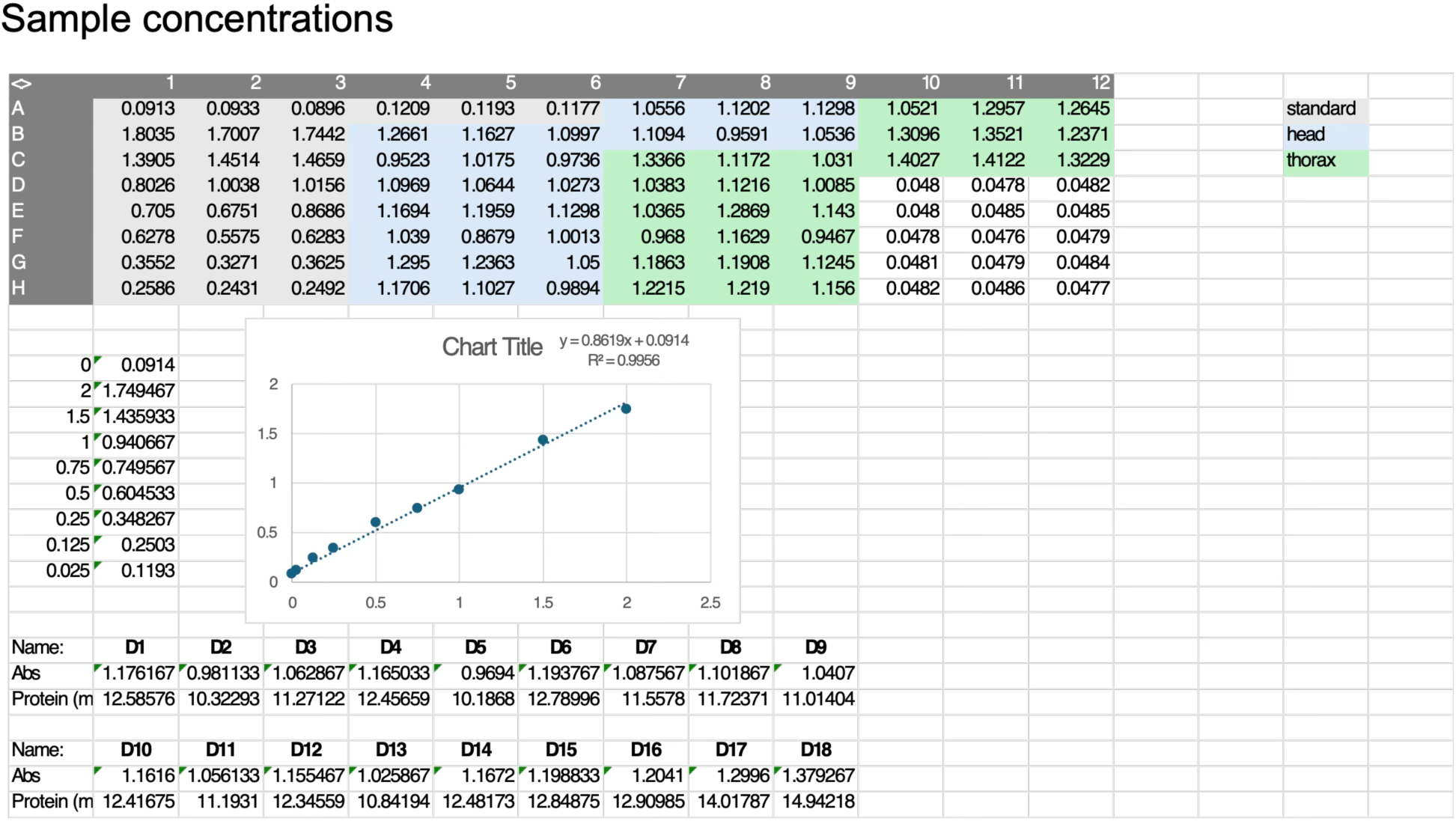
Proteomic sample concentrations.

**Figure S3:**
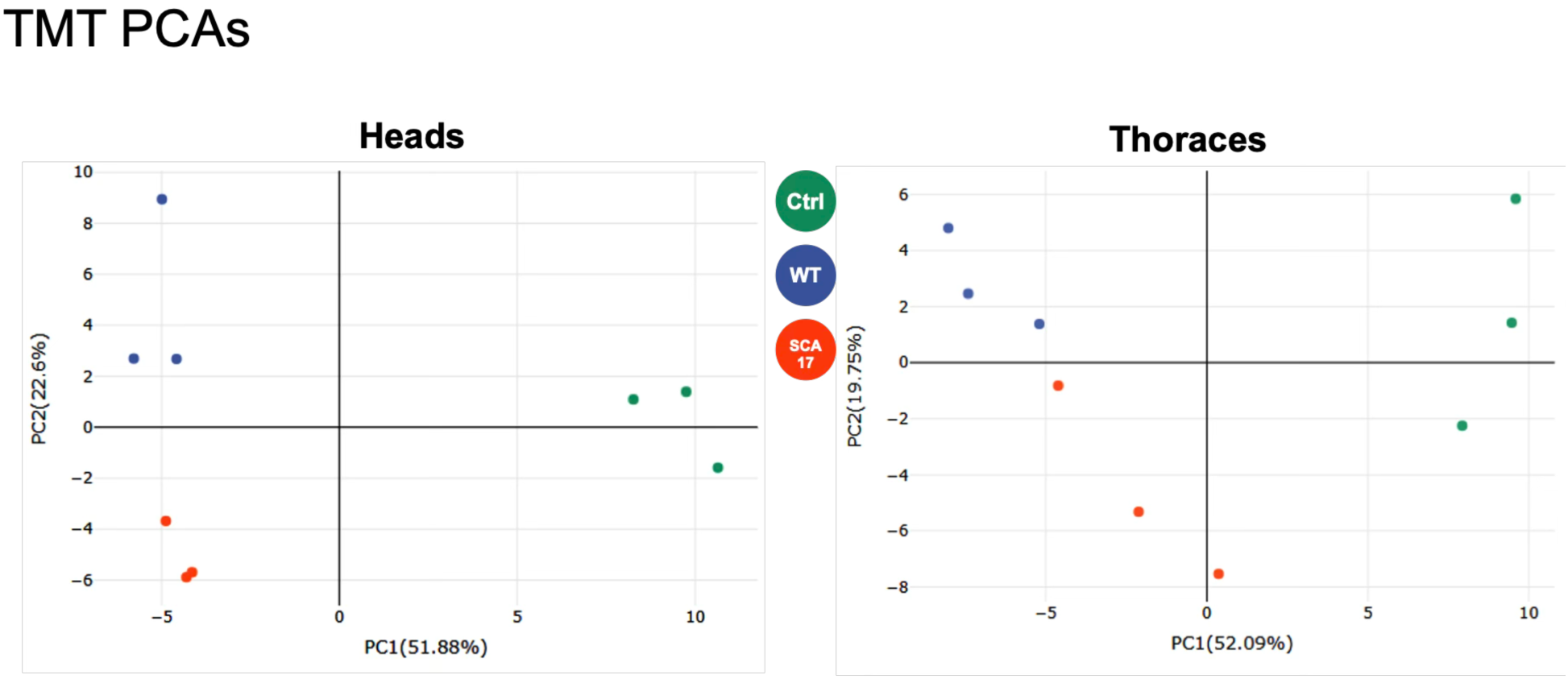
Principal component analyses for the proteomic samples showing heads (left) and thoraces (right).

**Figure S4:**
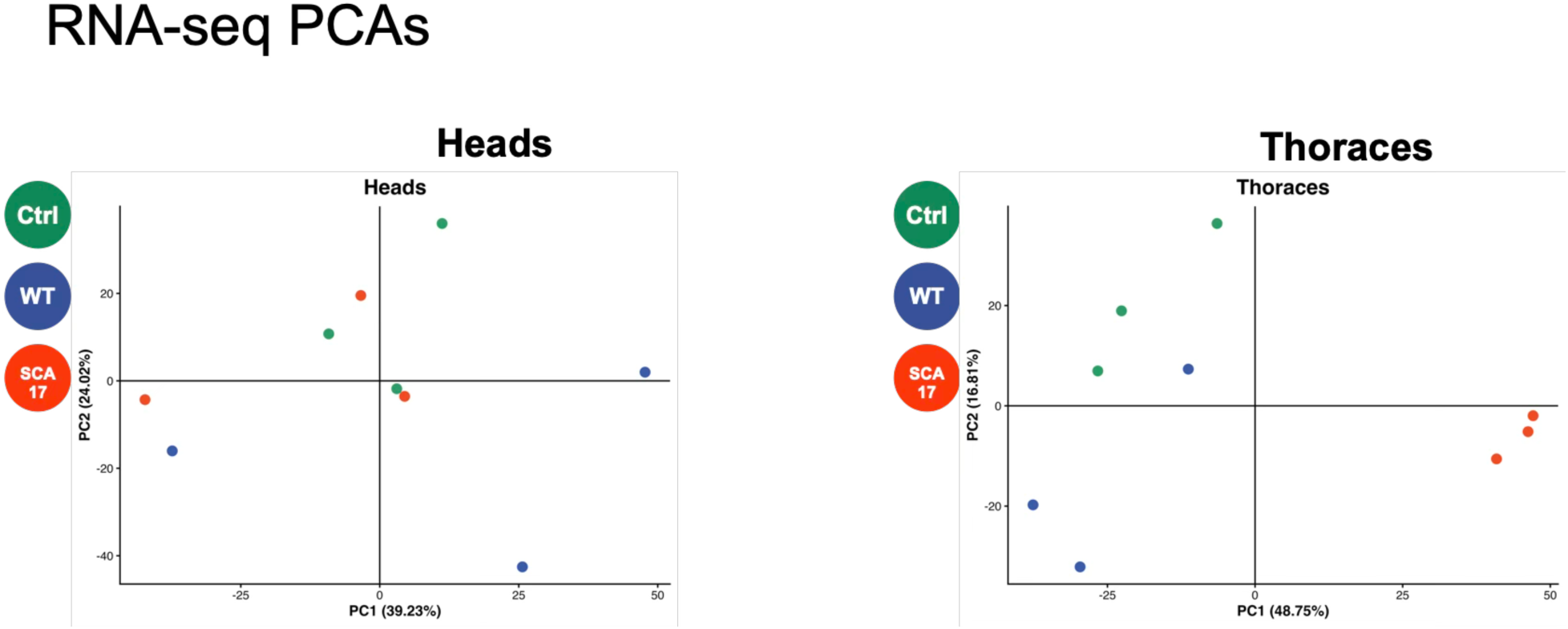
Principal component analyses for the RNA-seq samples showing heads (left) and thoraces (right).

